# Safety and uptake of fully oxidized β-carotene

**DOI:** 10.1101/2022.06.27.497750

**Authors:** Graham W. Burton, Trevor J. Mogg, Jacek Stupak, Felicity C. Stark, Susan M. Twine, Jianjun Li

**Author notes:** Corresponding author. Avivagen Inc., 100 Sussex Drive, Ottawa, Ontario, Canada K1A 0R6. *E-mail address:* (G.W. Burton). **Declaration of competing interest:** GWB and TJM are employees of Avivagen Inc. GWB owns shares in Avivagen Inc.

## Abstract

Spontaneous oxidation of β-carotene yields a polymer-rich product (OxBC) also containing small amounts of many apocarotenoids. OxBC extends β-carotene’s benefits beyond vitamin A, finding utility in supporting health in livestock, pets, and humans. Although naturally occurring OxBC is consumed in foods and feeds, a direct demonstration here of safety of synthetic OxBC supports its increasing usage. A toxicological study in rats showed a maximum tolerated single oral dose, an LD_50_, and a NOAEL of 5,000, 30,079 and 1875 mg/kg body weight, respectively. The repeat-dose 90-day oral toxicity study showed no adverse physiological or pathological effects. A first study of OxBC uptake by mice over 2-5 days into a select set of tissues showed OxBC already was naturally present. The highest levels were in liver, lung, and hamstring. Despite dosing, no net increases occurred in liver, kidney, lung, and muscle. Net increases occurred in urine, intestinal content, plasma, feces, spleen, and cecum, consistent with processing of OxBC and preferential elimination of polymer. Compared to the 4:1 polymer: apocarotenoid ratio of OxBC, polymer was enriched in liver and spleen and depleted in lung, kidney, hamstring, and abdominal muscle. The apparent control of OxBC in major tissues further supports its safety.

## 1. Introduction

β-Carotene spontaneously copolymerizes with ambient oxygen to form a naturally occurring substance found widely in plant-based foods (Burton et al., 2016; Burton et al., 2014; Schaub et al., 2017). The preponderance of polymer in oxidized β-carotene (OxBC) reflects the inherent preference of the highly unsaturated β-carotene backbone to add oxygen in a copolymerization process. There is significant evidence that the polymer-rich oxidized β-carotene extends β-carotene’s range of benefits beyond being a source of vitamin A (Johnston et al., 2014; Riley et al., 2021).

Synthetic OxBC is a complex mixture of compounds generated by the full, non-enzymatic air oxidation of pure β-carotene in solution. The spontaneous reaction generates two classes of compounds: 1) the newly recognized β-carotene-oxygen copolymer product (the “polymer”), and 2) a mixture of many low molecular weight apocarotenoid (“apoC”) breakdown products (Burton et al., 2014; Mogg and Burton, 2021). The apoC products are formed as by-products of the oxidative polymerization reaction. The polymer to apoC ratio is approximately 4:1 (w/w). The isolated polymer compound also has been shown to be partially susceptible to further breakdown into apocarotenoids under acidic and basic conditions (Mogg and Burton, 2021).

Synthetic OxBC is finding increasing use as a health-supporting product for livestock, pets, and humans. For example, livestock trials with low parts-per-million (ppm) supplementation of OxBC in feed have shown performance and health benefits over and above the benefits provided by vitamin and mineral premix supplements (Chen et al., 2020; Kang et al., 2018; McDougall, 2021; Riley et al., 2021). The absence of both β-carotene and vitamin A points directly to the involvement of the oxidation products as the source of OxBC’s beneficial effects.

With regard to safety, the negative outcomes of several human β-carotene intervention clinical trials (ATBC Cancer Prevention Study Group, 1994; Goodman et al., 2004; Omenn et al., 1994; Omenn et al., 1996a; Omenn et al., 1996b; Virtamo et al., 2014) drew attention to the possible involvement and potential toxicity of β-carotene oxidation compounds. We addressed this matter in a previous paper (Burton et al., 2021). To summarize, the physiological relevance of the cited supporting evidence, based entirely on *in vitro* model systems attempting to simulate oxidation conditions *in vivo*, was questioned in a review by an EFSA panel on the safety of β-carotene (European Food Safety Authority, 2012). Also, synthetic OxBC does not contain any of the long-chain, retinoid-like apocarotenoids (Burton et al., 2014) that have been suggested as potentially toxic agents that may adversely interfere with vitamin A retinoid receptor activity. OxBC’s apoC products are all non-retinoid, low molecular weight compounds with 8 to 18 carbon atoms, less than half of β-carotene’s 40 carbons and less than vitamin A’s 20. The two most abundant apocarotenoids are present at around 1% by weight. The oxidatively more reactive, long-chain apocarotenoid products (≥C_20_) that form early (Mordi et al., 1993) are ultimately consumed in the full β-carotene oxidation reaction and are therefore absent in OxBC. In plant food items these larger apocarotenoid compounds are present only in very low concentrations (Schaub et al., 2017). OxBC contains thirteen apocarotenoids (Mogg and Burton, 2021) that are designated as Generally Recognized As Safe (GRAS) human flavor agents (U.S. Food & Drug Administration).

OxBC is present naturally in feeds and foods. During storage or drying of plant products, β-carotene oxidation becomes significant (Burton et al., 2016), with the polymer being the main product (Schaub et al., 2017). Dietary intake of natural OxBC indirectly supports its safety. We have estimated dietary intake of natural OxBC from plant sources of β-carotene for humans and livestock (Burton et al., 2021). Vegetable powders and dried forages are rich sources of OxBC. The estimated exposure range for humans of 1-22 mg per serving is comparable to the recommended safe intake of β-carotene itself (<15 mg/d) (European Food Safety Authority, 2012). In livestock, OxBC in alfalfa can contribute ∼550-850 mg/head/d for dairy cattle, but in forage-deficient poultry feeds much less (∼1 ppm). Dairy cow intake of supplemental synthetic OxBC (300 mg/head/day) is comparable to OxBC that would be potentially available from traditional β-carotene-rich plant sources. Human intake of synthetic OxBC in meat from livestock fed OxBC is estimated to be similar to a single serving of food made with carrot powder.

The results of genotoxicity assays of synthetic OxBC have been reported (Burton et al., 2021). Although an Ames test showed weak-to-moderate mutagenicity at high concentrations of OxBC in only one cell line, a mouse micronucleus assay established an acute non-toxic dose of 1800 mg/kg body weight, and no bone marrow micronuclei were induced. The *in vivo* mouse results suggested that any potentially reactive compounds in OxBC are safely metabolized during acute exposure.

To substantiate the safety of chronic exposure to synthetic OxBC, we report here the results of a determination of the Maximum Tolerated Dose (MTD), the LD_50_, and the No Observed Adverse Effect Level (NOAEL) in 14-day and 90-day Repeat-Dose oral studies of OxBC in rats. Also, given the novelty of the polymer and the absence of any knowledge of its uptake from orally administered OxBC, we report here the results of the first study of the uptake of OxBC and its polymer and apoC fractions in mice.

## 2. Materials and methods

### 2.1. Oral toxicity studies in rats

Acute Oral Toxicity, Repeat-Dose 14-Day, and Repeat-Dose 90-Day Oral Toxicity studies were conducted by Anthem Biosciences Pvt. Ltd., Bangalore, India under contract to KGK Science, London, ON Canada. The care of animals complied with the regulations of the Committee for the Purpose of Control and Supervision of Experiments on Animals (CPCSEA) guidelines for laboratory animals published in the Gazette of India, 1998 and the Association for Assessment and Accreditation of Laboratory Animal Care International (AAALAC). The Form-B protocol for the conduct of the studies was reviewed and approved by the Institutional Animal Ethics Committee (IAEC Protocol No.: ABD/IAEC/PR/202-20-23; approved on June 1, 2020, and IAEC Protocol No.: ABD/IAEC/PR/251-21-24, Approved on: 06 August 2021). Guide for the Care and Use of Laboratory Animals, National Research Council, 2011.

#### 2.1.1. OxBC test article

OxBC was prepared commercially by Allied Biotech Corp., Taipei, by spontaneous air oxidation of pure synthetic β-carotene in ethyl acetate solution, essentially based on the laboratory procedure (Burton et al., 2014), and stored in a freezer when not in use.

#### 2.1.2. Acute oral toxicity study

The study was conducted to determine the acute systemic toxicity potential and MTD of OxBC in female Sprague Dawley rats. Animals were acclimatized for a minimum period of 5 days prior to OxBC administration. A total of 15 animals were used with three females for each level of OxBC tested. Formulations of OxBC were prepared in a vehicle comprising DMSO:PEG 400:propylene glycol (1:2:2 by volume). The dose volume was 10 mL/kg body weight.

Overnight fasted animals were administered a single dose or a divided dose within 24 hours by gavage followed by an observation period of 14 days to select the dose levels for a subsequent repeat-dose 14-day toxicity study. OxBC was dosed at 5,000 mg/kg, 10,000 mg/kg (5000 mg/kg b.i.d.), 2,500 mg/kg, 7,500 mg/kg (3750 mg/kg b.i.d.) and 1,250 mg/kg body weight, respectively. After dosing all animals were observed at 20 to 30 min, 1 h ±10 min, 2 h ±10 min and 4 h ±10 min post dose, and they were then observed once daily for clinical signs and twice daily for mortality or morbidity for 14 days. Non-fasted body weights were recorded on day 0 (last day of acclimatization) for all animals; day 8, day 14 and day 15 for surviving animals and before necropsy for sacrificed animals found dead or moribund. On day 15, all surviving animals were euthanized with an overdose of carbon dioxide and subjected to gross necropsy examination. Dead and euthanized moribund animals were subjected to necropsy at the earliest opportunity.

#### 2.1.3. Repeat-dose 14-day oral toxicity study

The study was conducted to determine the systemic toxic potential of OxBC upon repeated once-daily administration for 14 consecutive days by gavage in Sprague Dawley rats and to determine the NOAEL.

Twenty male and twenty female animals were assigned to four groups of five animals per gender, viz., vehicle control, low-dose, mid-dose and high-dose. Formulations were prepared in a DMSO:PEG 400:propylene glycol vehicle (1:2:2 by volume). The control group was administered vehicle alone orally. The OxBC formulations were administered by gavage at dose levels of 1,250 (low-dose), 2,500 (mid-dose) and 5,000 (high-dose) mg/kg body weight once daily for 14 consecutive days. The dose volume was 10 mL/kg body weight. Animals were observed once daily for clinical signs, twice daily for mortality or morbidity and once weekly for detailed clinical examination. Body weights and feed consumption were recorded at weekly intervals. At the end of the experimental period (Day 15), blood samples were analyzed for hematology parameters. Harvested plasma specimens were analyzed for coagulation and clinical chemistry parameters. Subsequently, the animals were euthanized and subjected to gross pathological examination. Specified organs were collected, weighed, and preserved in a suitable fixative for histopathological evaluation.

#### 2.1.4. Repeat-dose 90-day oral toxicity study

A repeat dose 90-day toxicity study was performed to determine the systemic toxicity potential of OxBC when administered by gavage at graded dose levels administered once daily for a period of 90 consecutive days in Sprague Dawley rats. The data from the study allowed for the characterization of OxBC toxicity, a dose response relationship, if any, and the determination of the NOAEL.

Study Design. Fifty males and fifty females were assigned to 4 groups of ten animals of each gender per group for the main study groups, and two recovery groups of five animals of each gender per group. Randomization was based on body weights on the last day of acclimatization, with body weight variation of animals not exceeding ± 20% of the mean body weight of each gender. Animals were acclimatized six days for males and seven days for females. The animals were orally administered once daily with vehicle alone or OxBC for 90 consecutive days at a dose volume of 5 mL/kg body weight at dose levels of 625, 1,250 and 1,875 mg/kg body weight, designated as low-, mid-, and high-dose, respectively. The doses were based on the 14-day repeated dose toxicity study. The groups were: G1, control (vehicle); G2, low-dose; G3, mid-dose; G4, high-dose; G1R, control recovery; G4R, high-dose recovery. The recovery period was 28 days.

The OxBC test item was formulated in DMSO: PEG 400: propylene glycol (1:2:2) vehicle, as used for the prior acute maximum tolerated dose and the 14-day repeat-dose studies. The formulation was prepared fresh prior to administration. The final pH and appearance of the formulation were recorded on Day 1 and the last week of dosing. The stability and homogeneity of the dose formulations were performed in Week 1 and during Weeks 11, 12. The acceptance criteria were ± 15% of the nominal concentration of geronic acid, a marker compound of OxBC. A sample of the vehicle control group also was analyzed for geronic acid to rule out any possible contamination with OxBC.

During the period of administration, the animals were observed closely each day for signs of toxicity. Any animals that died or were euthanized during the treatment period were necropsied. At the end of treatment, the surviving animals were euthanized and necropsied. Mortality and morbidity were monitored twice daily during the study. All animals were observed once daily for visible clinical signs.

Detailed clinical examinations were performed during the acclimatization period and weekly during the treatment period. Animals were examined for, but not limited to, changes in skin, fur, eyes, mucous membranes, occurrence of secretions, excretions, and autonomic activity (e.g., lacrimation, piloerection, unusual respiratory pattern), changes in gait, posture, and response to handling, as well as the presence of clonic or tonic movements. Body weights and feed consumption were recorded weekly. Ophthalmological examination was performed on all the animals before start of treatment and when on vehicle (G1) and in the high-dose treatment (G4) groups at the end of the study (Week 13). Functional observation tests were carried out during Weeks 12, 13 for groups G1 and G4 and during Week 17 for recovery groups G1R and G4R. Urinalysis was performed on all groups in Week 13 and the G1R and G4R groups in week 17. Hematology, coagulation, clinical chemistry, and hormonal analyses were performed on blood collected from overnight-fasted animals of all groups on Day 91 and the G1R and G4R groups on Day 119. Terminal vaginal cytological examination of all females was performed one day prior to necropsy and the estrus cycle was determined. Gross necropsy was conducted on the animals of groups G1-G4 sacrificed at Day 91 and on groups G1R and G4R sacrificed at Day 119. Tissues were collected and organs weighed. Organs from the G1 and G4 groups were preserved in suitable fixative for histopathological examinations.

### 2.2. Mouse OxBC uptake study

#### 2.2.1. Compounds

OxBC, prepared as described earlier (Burton et al., 2014), was used to prepare the OxBC polymer fraction and the low molecular weight OxBC apoC fraction.

OxBC polymer. OxBC (2.02 g) was dissolved in ethyl acetate (5 mL) in a 100 mL round bottom flask with stir bar. Hexane (50 mL) was added dropwise with stirring and after 1 hour the liquid was decanted. The residue was rinsed with hexane (3 × 3 mL), blown dry with N_2_, redissolved in ethyl acetate and concentrated on the rotary evaporator at < 10 torr, 40°C to give a crisp, yellow, sponge-like solid. The precipitation was repeated four more times and the final product dried under vacuum to give the OxBC polymer (1.04 g) as a yellow, sponge-like solid.

OxBC apoC was prepared as has been described briefly earlier (Burton et al., 2016; Burton et al., 2014). OxBC (250 mg) was placed in a glass centrifuge tube (15 mL) and dissolved in ethyl acetate (1.2 mL). OxBC polymer was precipitated from the stirred solution by adding hexane (12 mL) dropwise. The mixture was centrifuged (10 min), the supernatant transferred to a round bottom flask (50 mL), and the liquid removed by rotary evaporation. The residual yellow oil was dried under vacuum to give the crude apoC fraction (130 mg), which was purified by further hexane precipitations from concentrated ethyl acetate solutions as follows: a sample (95 mg) was placed in a glass centrifuge tube (15 mL), dissolved in ethyl acetate (0.4 mL), and hexane (12 mL) added dropwise with stirring. The stir bar was removed, the sample centrifuged (10 min) and the supernatant transferred to a round bottom flask and concentrated with rotary evaporation. After drying under vacuum, the residual oil (65.4 mg) was subject to a repeat purification step by dissolving the oil in ethyl acetate (0.2 mL) in a round bottom flask with stirring and adding hexane (20 mL) dropwise. The resulting cloudy liquid was drawn into a syringe and passed through a Teflon syringe filter (0.2 µm). The flask and syringe were rinsed with hexane and the rinsings passed through the same filter. All filtrates were combined, concentrated by rotary evaporation and dried under vacuum to give pure OxBC apoC (54 mg).

Two deuterium-labelled standards, d_12_-OxBC polymer and d_6_-dihydroactinidiolide (d_6_-DHA), were used for LC-MS determination of OxBC polymer and OxBC apoC concentrations, respectively. d_12_-OxBC was synthesized from 16,17,16’,17’-[(C^2^H_3_)_4_]-dodecadeuterium-labelled β-carotene by the same method used to prepare unlabelled OxBC (Burton et al., 2014). The solid d_12_-OxBC polymer compound was obtained by successive solvent precipitations from d_12_-OxBC, as described above for the unlabelled compound. Details of the syntheses of d_12_-β-carotene, d_12_-OxBC and the d_12_-polymer will be reported elsewhere. The synthesis of d_6_-DHA has been described (Burton et al., 2014).

All compounds were stored in a freezer until required.

#### 2.2.2. Animals

Twelve female BALB/c mice, aged 6-8 weeks and weighing approximately 18 g, were purchased from Charles River Laboratories. The mice were maintained at the National Research Council of Canada (NRC) in accordance with the guidelines of the Canadian Council on Animal Care. All procedures performed on animals used in the study were in accordance with regulations and guidelines reviewed and approved by the NRC Human Health Therapeutics Ottawa Animal Care Committee (Protocol # 2020.01). For the first 6-8 weeks of their life the mice were fed the Charles River Rat and Mouse 18% (Auto) diet, 5L79* (LabDiet, Quakertown, PA). During the period the mice were housed at the NRC facility, from November 18, 2020, until study end on December 4, 2020, the animals were fed the 2014 Teklad Global 14% Protein Rodent Maintenance Diet (Envigo, Indianapolis, IN). At the NRC the mice had ready access to ultrapure water.

#### 2.2.3. Protocol

OxBC, a highly viscous, water-insoluble liquid, was formulated in aqueous 30% DMSO (v/v). A stock solution of pure OxBC (200 mg/mL) was prepared in sterile DMSO. OxBC solution and sterile water (milliQ) were separately aliquoted into 15 mL polypropylene tubes in amounts required for use each day and stored frozen. On the days of dosing the individual frozen OxBC aliquots and water aliquots were thawed, mixed, and used within 4 hours. No OxBC precipitated out of solution. The vehicle solution was prepared similarly using DMSO in place of OxBC stock solution. Mice received either OxBC (300 mg/kg body weight) or the DMSO vehicle daily by gavage for 2 days or 5 days. Each treated mouse (20 g) received 100 µL of a solution of OxBC in DMSO (30 µL) and water (70 µL), providing 6 mg of OxBC. Control mice received the vehicle of DMSO (30 µL) and water (70 µL).

Mice were assigned randomly into 4 groups of 3 mice each. Two groups were treated orally once daily with OxBC (100 µL) for 2 days and 5 days, respectively. The two control groups were treated orally with vehicle (100 µL) for 2 days and 5 days, respectively.

The mice were gavaged with OxBC or vehicle each day at 8 am. On Day 1, blood was collected from all mice at least one hour after gavage. On Days 2 and 5, blood, urine and feces were collected 1 hour after gavage from the 2-day and 5-day OxBC-dosed and vehicle-dosed groups, respectively, followed by euthanasia and tissue collections. Vehicle-treated mice were euthanized first. Tissues and tissue contents were harvested in the following order: 1) urine, feces; 2) blood (plasma); 3) liver (all lobes), lung (5 lobes), leg muscle (hamstrings, 2 legs), abdominal muscle, spleen; 4) whole stomach, intestinal flush (10 mL), flushed small intestine, cecum, large intestine.

Blood (100-200 µL) was collected by cheek vein puncture into heparin coated BD microtainer^®^ blood collection tubes (Becton Dickinson, Franklin Lakes, NJ), then inverted to mix and immediately placed on wet ice.

Urine was collected into micro-centrifuge collection tubes by pressing the tube against the abdomen of the mouse. The collected urine was placed on dry ice before being transferred for storage at −80°C. The mouse was then placed in a wide mouth jar with lid and 3-4 fecal pellets were collected into a micro-centrifuge tube and placed on wet ice before transfer to storage at −80°C.

Mice were euthanized with isoflurane (4%) and oxygen (2%). Tissues were collected into petri dishes, cut as described below, blotted dry, weighed, and transferred to labelled homogenization tubes containing cold PBS (2 mL) and placed on wet ice. The tissues were collected as follows: the first layer of abdominal muscle was cut out and minced with scissors, the whole spleen was excised, and the fat trimmed off, both kidneys were collected and cut in half across the narrower middle. All liver lobes were collected with the fat trimmed off.

Hamstrings were taken and cut into small sections. All 5 lung lobes were collected and cut into small pieces. The stomach and attached intestine were removed from the abdominal cavity. The stomach was cut off from the small intestine and moved to a fresh petri dish where it was cut open along one side and inverted. The stomach was flushed with PBS (10 mL) to remove large debris, then placed in a 15 mL falcon tube with fresh PBS, the tube was capped and shaken to wash the tissue and loosen more debris from the tissue. The stomach was further washed in multiple falcon tubes containing fresh PBS until no more debris was observed in the wash solution. Finally, the stomach was cut into 4 sections and placed in a homogenization tube. The intestine was spread out, breaking all the connective tissue, and removing any fat. The intestinal tube was transferred to a fresh petri dish and flushed with PBS (10 mL) that was collected into a 15 mL falcon tube and placed on ice. The intestine was cut into small intestine, cecum and large intestine segments that were removed, cut open, washed in PBS, cut into 3-4 pieces, and transferred to homogenization tubes. All samples were kept on wet ice throughout collection and processing, except for urine, which was placed in dry ice.

Homogenization tubes containing weighed tissue samples were transferred from ice to a Precellys Evolution homogenizer (Bertin Technologies SAS, Montigny-le-Bretonneux, France). The samples were dissociated using three 10 second pulses with 30 second intervals and then placed on ice before aliquoting the homogenate into three 1.5 mL microcentrifuge tubes for storage at −80°C. Total sample volumes were 2 mL, except for liver (3 mL).

#### 2.2.4. LC-MS/MS analytical method development

Supplementary Material *S1* describes in detail the LC-MS/MS method developed to analyze OxBC’s two principal components, the major OxBC polymer fraction and the minor apoC fraction, in the determination of their levels in animal tissues and fluids.

LC-MS analysis of the OxBC apoC fraction confirmed the presence of a multitude of apocarotenoids. LC-multiple reaction monitoring (MRM), the method of choice for sensitive and selective quantitative analysis of small molecules, yielded a greatly simplified chromatogram containing four of the most abundant OxBC apocarotenoid compounds identified previously (Burton et al., 2014), namely dihydroactinidiolide (DHA), ß-ionone, ß-ionone-5,6-epoxide and geronic acid. The same pattern of detected compounds was obtained for plasma spiked with OxBC apoC.

The OxBC polymer was not amenable to direct analysis by mass spectrometry. However, NaOH treatment of the polymer generated the same four apocarotenoids seen in the LC-MRM analysis of OxBC apoC. Successful indirect detection of OxBC polymer in this manner in spiked muscle and liver homogenates and plasma confirmed NaOH treatment afforded a viable approach to the determination of OxBC polymer in tissues and body fluids.

#### 2.2.5. LC-MS/MS analysis of OxBC in tissues and body fluids

Calibrations and quantitation of the OxBC polymer fraction and the apoC fraction were carried out using the deuterium-labelled internal standards, d_12_-OxBC polymer and d_6_-DHA, respectively. Both standards were added together to tissue homogenates and body fluids.

Calibration of OxBC polymer was performed by adding a specially prepared internal standard, NaOH-treated d_12_-OxBC polymer, to a series of dilutions of NaOH-treated OxBC polymer samples in methanol. NaOH digestion of unlabelled OxBC polymer and d_12_-OxBC polymer released apocarotenoid marker compounds for LC-MRM analysis.

The NaOH-treated d_12_-OxBC polymer internal standard was prepared by adding aqueous NaOH (1M, 240µL) to OxBC polymer (10 mg) in methanol (1 mL) and heating the solution at 75°C for 4 h with shaking (1100 rpm). After cooling, the solution was brought to pH∼5 with aqueous HCl (4M, 60 µL) to give a final concentration of 7.69 µg/µL OxBC polymer NaOH digest. A d_12_-OxBC polymer digest was prepared similarly, as follows. A solution of d_12_-OxBC polymer (10 µL, 100 µg), methanol (65 µL) and aqueous NaOH (1M,18 µL) was heated at 75°C for 4 h with shaking (1100 rpm). After cooling, the solution was neutralized with HCl (4M, 4.5 µL) and brought to a final volume of 100 µL with methanol to give a final concentration of 1 µg/µL of the d_12_-OxBC polymer NaOH digest.

Calibration was performed using six solutions of OxBC polymer digest, providing LC-MS on-column amounts ranging from 0.5 ng to 50 µg with 20 ng of d_12_-OxBC polymer NaOH digest internal standard.

Calibration of OxBC apoC was carried out by adding the d_6_-DHA internal standard to a series of six dilutions of OxBC in methanol that provided an LC-MS on-column range of 0.5 ng to 50 µg OxBC with 0.8 ng d_6_-DHA. The calibration of the DHA present in the apoC fraction of OxBC versus OxBC itself provided an indirect calibration against the OxBC apoC fraction knowing that the apoC fraction is 20% of OxBC by weight.

Quality control (QC) samples and tissue and body fluid samples were processed by extracting with hexane to remove the OxBC apoC compounds and added d_6_-DHA. The hexane fraction containing the apoC fraction and d_6_-DHA was evaporated and analyzed for OxBC apoC compounds. The remaining aqueous solution was extracted with ethyl acetate to recover OxBC polymer and d_12_-OxBC polymer, which were digested with NaOH for indirect quantitation of the parent compound by LC-MRM analysis.

The general procedure was as follows: to tissue homogenates (500 µL), body fluid samples (20-500µL), or methanol solutions (25 µL) of the low-QC (5 µg OxBC) and high-QC (25 µg OxBC) samples, were added 2 µL of a methanol solution of d_12_-OxBC polymer (1 µg) and d_6_-DHA (40 ng). Each sample was vortex mixed for 1 min with ethanol (0.4 mL). Hexane (0.4 mL) was added, vortex mixed for 1 min, then water (250 µL) was added, and the mixture was centrifuged for 2 min at 13,000 rpm. The top hexane layer containing the apoC fraction was removed and put aside. The hexane extraction process was repeated 3 times for a total of 4 extractions. The hexane washes were combined, the solvent removed using a vacuum centrifuge concentrator, and the residue taken up in methanol (50 µL) for LC-MS analysis. The remaining aqueous ethanol fraction was split between two vials. Ethyl acetate (250 µL) and water (500 µL) were added to each vial, vortex mixed for 1 min and then centrifuged at 13,000 rpm for 2 min. The ethyl acetate layer containing OxBC polymer was removed. The extraction process was repeated after adding ethyl acetate (250 µL) to the remaining aqueous fraction. The combined ethyl acetate fractions were taken to dryness using vacuum centrifugation over ∼2 h. The resultant pellet was dissolved in ethyl acetate (25 µL) with vortex mixing and sonication. Hexane (1 mL) was added, and the mixture was placed in a refrigerator for 30 min to precipitate the OxBC polymer. The precipitate was spun down and the solvent removed and discarded. The precipitate, dissolved in methanol (100 µL), was heated with NaOH at 75°C for 4 h, acidified to pH 4 with HCl (4M, 6 µL) to give a final volume of 130 µL of isolated OxBC polymer. All solutions were stored in a freezer pending LC-MS analysis.

For the mouse study, LC-MRM was performed on a TSQ Quantiva triple-stage quadrupole mass spectrometer coupled to a Dionex Ultimate 3000 micro HPLC system (Thermo Fisher Scientific Inc, Waltham, MA, USA) using electrospray ionization. The MRM method was employed to detect and identify low molecular weight apocarotenoids present in the isolated apoC fraction and released in the NaOH-treated polymer fraction. DHA was chosen as the marker compound for both the polymer and apoC fractions.

## 3. Results

### 3.1. Oral toxicity studies in rats

#### 3.1.1. Acute oral toxicity study

Rats dosed 5,000 mg/kg body weight OxBC showed clinical signs of excessive salivation, polyuria and perineum wetting. A sweet aroma in the cage was observed. All animals were found to be normal at 4 h post dose. In animals dosed 5,000 mg/kg body weight b.i.d for a total daily administration of 10,000 mg/kg/day body weight there were clinical signs of excess salivation, dark yellowish urination, a sweet aroma in the cage, hypoactivity, piloerection, dehydration, abdominal breathing, hunched posture, and somnolence. One animal was found dead.

Animals administered 2,500 mg/kg body weight showed clinical signs of mild salivation at 30 min post-dose and were normal out to Day 15. Animals dosed 3,750 mg/kg body weight b.i.d for a total daily administration of 7,500 mg/kg/day body weight showed clinical signs of excess salivation, hypoactivity, dehydration, and tremor. One animal was found dead post-dose at 30 min after the second dose. The other two animals were normal at Day 2 post-dose. Animals administered 1,250 mg/kg body weight were normal out to Day 15.

After administration at the specified dose levels no adverse effects on body weight were observed in the animals that survived out to the scheduled sacrifice endpoint (Day 15). Gross necropsy examination of the surviving animals at the terminal sacrifice, as well as deceased animals, revealed no abnormal gross pathological findings.

Under the experimental conditions and doses employed, the maximum tolerated dose was determined to be 5,000 mg/kg body weight. The calculated single dose LD_50_ of OxBC by probit analysis was 30,079 mg/kg body weight.

#### 3.1.2. Repeat dose 14-day oral toxicity study

The NOAEL of OxBC administered once daily for 14 consecutive days by oral gavage to Sprague Dawley rats for both genders was determined to be 1250 mg/kg body weight. The following clinical observations were made in comparison to the vehicle control group:

Clinical signs and mortality or morbidity. No mortality or morbidity was observed in the vehicle control and low dose groups for both genders, and in mid-dose males. One female in the mid-dose group (2,500 mg/kg) found moribund was euthanized. Two males and one female were found dead in the high-dose group (5,000 mg/kg). Clinical signs of hypoactivity and dehydration were observed in the mid- and high-dose group for both genders. On detailed clinical examination, hypoactivity, dehydration, sunken eyeballs, and, in the moribund animals, abdominal breathing and distended abdomens were observed in mid- and high-dose groups, with varying incidences in both genders.

Weight and body weight gain. There were no adverse effects on body weight and body weight gain in the low-dose group for both genders. Statistically significant decreases in body weight and body weight gain were observed in high-dose males. Although statistically significant changes were not observed in mid-dose males and females and high-dose females, a trend toward decreased body weight and body weight gain was observed in these specific groups.

Feed consumption. There were no adverse effects on feed consumption in the low-dose group for both genders compared to the vehicle control group. In high-dose males, a decrease in Week 1 feed consumption was observed with a statistically significant decrease in Week 2.

Clinical pathology, hematology, coagulation. There were no adverse effects on hematology or coagulation parameters in all treatment groups for both genders.

Clinical chemistry. No adverse effects on clinical chemistry parameters were observed in all treatment groups for both genders, although dose-dependent decreases in triglycerides were observed in males, which may have been secondarily related to decreases in body weight.

Gross necropsy. No gross pathological findings were observed in either gender in all treatment groups.

Organ weights. No adverse effects were observed on the absolute and relative organ weights of all treatment groups for both genders.

Histopathology. No treatment-related histopathological findings were observed in any of the specified organs evaluated in the high-dose groups.

#### 3.1.3. Repeat dose 90-day oral toxicity study

The NOAEL for OxBC administered once daily for 90 days was determined to be 1875 mg/kg body weight. No adverse effects were observed in any of the treatment groups. The following clinical observations were made in comparison to the vehicle control group:

Mortality and morbidity, clinical signs, and detailed clinical examination. No mortality or morbidity and no adverse clinical signs were observed in all the treatment groups. All the animals survived until the scheduled sacrifice.

Ophthalmological examination. No treatment-related ophthalmological abnormalities were observed in the animals of the high dose group for both genders compared to the vehicle control group.

Body weight, body weight gain and feed consumption. No treatment-related adverse effects on body weight and body weight gain were observed in all the treatment groups for both genders compared to the respective vehicle control groups.

Functional observation battery tests and neurological examination. Functional observation battery tests revealed no treatment-related adverse effects on the parameters examined in the high dose group compared to the vehicle control group.

Vaginal cytology. Treatment-related adverse effects were not observed on the estrus cycle in females as the stages of estrus cycle were evenly distributed among all the groups.

##### Clinical Pathology

Urinalysis. No treatment-related adverse effects were observed in the group mean values or in incidences of semi-quantitative observations in any of the treatment groups of both genders compared to the vehicle control group.

Hematology, coagulation, clinical chemistry and hormonal parameters. No treatment-related adverse effects on hematology, coagulation, clinical chemistry and hormonal parameters analyzed (T3, T4 and TSH) were observed in any of the treatment groups of both genders compared to the vehicle control group.

Gross necropsy. No treatment-related gross pathological findings were observed in any of the treatment groups of both genders.

Organ weights. No treatment-related adverse effects were observed on the absolute and relative organ weights of any of the treatment groups of both genders compared to the vehicle control group.

Histopathology. Microscopic examination of the collected tissues showed no treatment-related histopathological findings in the high dose treatment group of both the genders compared to the vehicle control group. The lower dose of the main study and recovery groups were not evaluated.

### 3.2. OxBC mouse uptake study

#### 3.2.1. LC-MRM analytical method

As reported in Supplementary Material *S1*, LC-MS analysis of the OxBC apoC fraction showed the presence of a multitude of apocarotenoids. LC-MRM analysis yielded a greatly simplified chromatogram. Three major OxBC apocarotenoid peaks were clearly apparent, corresponding to compounds identified previously as the most abundant in the OxBC apoC fraction (Burton et al., 2014), namely, DHA, ß-ionone and ß-ionone-5,6-epoxide. A small peak corresponding to geronic acid also was present. LC-MRM analysis of a methanol extract of vacuum-dried OxBC apoC-spiked serum showed an almost identical pattern for the same peaks.

The OxBC polymer, however, was not directly detectable by mass spectrometry (Supplementary Material *S1*, section 3.2). Previously, NaOH digestion of the polymer had been shown to release apocarotenoids (Mogg and Burton, 2021). Application of NaOH treatment of the polymer coupled with LC-MRM analysis revealed the presence of the same four apocarotenoids seen in the LC-MRM analysis of OxBC apoC. Use of the NaOH/LC-MRM combination confirmed recovery of OxBC polymer spiked into plasma and into homogenates of muscle and liver. These results supported the application of the NaOH/LC-MRM approach for the detection and estimation of levels of OxBC polymer in mouse tissues and body fluids. LC-MRM analysis also was used for direct estimation of OxBC apoC in hexane extracts of tissues, plasma, and serum.

Because LC-MRM analysis subsequently showed that only DHA was consistently present in all tissues and fluids of OxBC-dosed mice, it was used as the marker compound for estimating OxBC polymer and apoC levels. In principle, these measurements made it possible to estimate in each tissue and fluid the total OxBC, i.e., the sum of the polymer and apoC fractions, and the ratio of the polymer fraction to the apoC fraction.

Calibrations. With an emphasis on the OxBC polymer, the main component of OxBC, linear calibrations were determined for each of the two main DHA transitions, 181.1→107.1 and 181.1→135.1 amu. Both transitions gave a linear response, with R^2^ values higher that 0.998 over a 1000-fold range of concentration. Three independent calibration curves for each transition were run to assess reproducibility. Relative standard deviations (RSD) of peak areas were found to be acceptable, with the majority under 13% and the highest outlier at 28%. For both DHA transitions the limit of detection (LOD) was 0.08 ng/µL, the lower limit of quantification (LLOQ) was 0.25 ng/µL, and the upper limit of quantification (ULOQ) was 10 µg/µL. The inter-day reproducibility of three independent calibrations prepared over a period of several weeks was evaluated. For the 181.1→107.1 and 181.1→135.1 transitions the RSD values for slope were 4.06% and 2.76%, respectively. The corresponding RSD values for R^2^ were 0.135% and 0.038%. The accuracy from 12 independent inter-day measurements of the low-QC (50 ng/µL) was 74.8% (RSD 38.1%) and of the high-QC (250 ng/µL) was 71.7% (RSD 36.5%).

For the OxBC apoC fraction the accuracy of the low-QC (50 ng/µL) was 105.2% (RSD 26.3%) and of the high-QC 70.1% (RSD 39.8%).

#### 3.2.2. OxBC polymer and apoC content of tissues and fluids

Tables 1 and 2 in Supplementary Material *S2* present OxBC polymer and OxBC apoC data, respectively, for individual tissue and body fluids of individual control and OxBC-dosed mice after 1 h dosing (plasma only) and after 2 days and 5 days daily dosing (plasma and tissues). Both OxBC polymer and OxBC apoC forms were found to be present in all tissues and fluids of control mice. In the dosed mice the between-mice variability of the values obscured any potential trends in OxBC tissue and fluid content occurring with duration of OxBC dosing. Therefore, in Table 1 the data for Days 2 and 5 are combined and presented in the form of total OxBC, i.e., polymer + apoC, and the ratio of polymer to apoC, i.e., polymer/apoC.

**Table 1.**
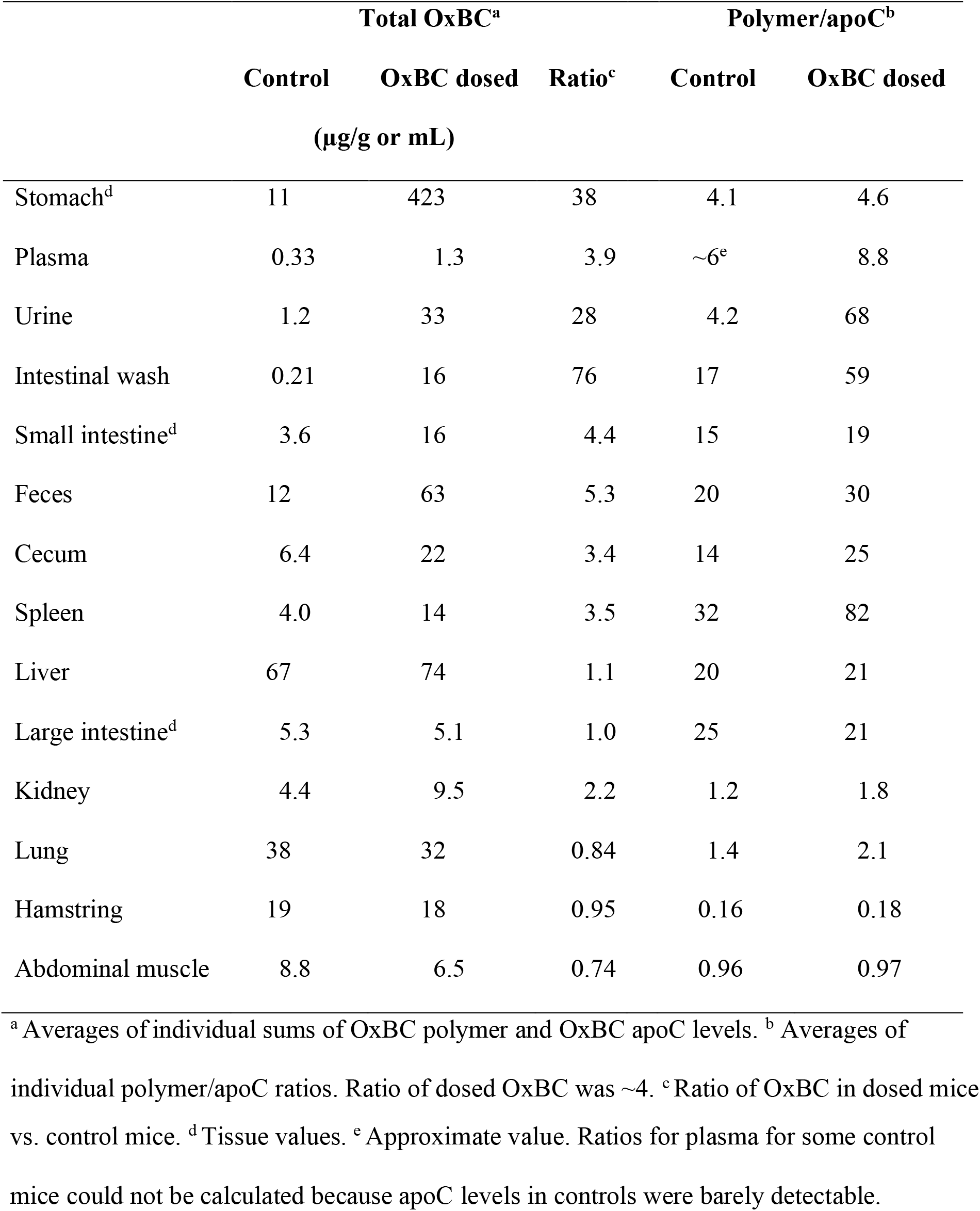
Average concentrations of OxBC and ratios of OxBC polymer to OxBC apocarotenoids in tissues and fluids of mice dosed orally with OxBC (300 mg/kg body weight) daily for 2 and 5 days vs. control mice dosed with the same volume of 30% aqueous DMSO vehicle alone.

|  | Total OxBC<sup>a</sup> |  |  | Polymer/apoC<sup>b</sup> |  |
| --- | --- | --- | --- | --- | --- |
|  | Control | OxBC dosed | Ratio<sup>c</sup> | Control | OxBC dosed |
|  | (µg/g or mL) |  |  |  |  |
| Stomach<sup>d</sup> | 11 | 423 | 38 | 4.1 | 4.6 |
| Plasma | 0.33 | 1.3 | 3.9 | ~6<sup>e</sup> | 8.8 |
| Urine | 1.2 | 33 | 28 | 4.2 | 68 |
| Intestinal wash | 0.21 | 16 | 76 | 17 | 59 |
| Small intestine<sup>d</sup> | 3.6 | 16 | 4.4 | 15 | 19 |
| Feces | 12 | 63 | 5.3 | 20 | 30 |
| Cecum | 6.4 | 22 | 3.4 | 14 | 25 |
| Spleen | 4.0 | 14 | 3.5 | 32 | 82 |
| Liver | 67 | 74 | 1.1 | 20 | 21 |
| Large intestine<sup>d</sup> | 5.3 | 5.1 | 1.0 | 25 | 21 |
| Kidney | 4.4 | 9.5 | 2.2 | 1.2 | 1.8 |
| Lung | 38 | 32 | 0.84 | 1.4 | 2.1 |
| Hamstring | 19 | 18 | 0.95 | 0.16 | 0.18 |
| Abdominal muscle | 8.8 | 6.5 | 0.74 | 0.96 | 0.97 |

<sup>a</sup> Averages of individual sums of OxBC polymer and OxBC apoC levels. <sup>b</sup> Averages of individual polymer/apoC ratios. Ratio of dosed OxBC was ~4. <sup>c</sup> Ratio of OxBC in dosed mice
vs. control mice. <sup>d</sup>Tissue values. <sup>e</sup> Approximate value. Ratios for plasma for some control mice could not be calculated because apoC levels in controls were barely detectable.

Although the small number of mice and the between-mice variability precluded any statistical comparisons, gross trends were apparent. The results divided into two groups, depending on how the tissues and fluids responded to dosing.

Tissues and fluids in the first group, comprising stomach, small intestine, intestinal wash, plasma, urine, feces, cecum, and spleen, showed a substantial excess of OxBC in the OxBC-dosed mice compared to controls. The dosed mice also showed a substantial excess of polymer over apoC relative to the 4:1 polymer/apoC ratio of the dosed OxBC, with one exception. In stomach tissue, which for dosed mice had the highest content of OxBC of all tissues examined, the ratio was ∼4, essentially the same as the initial OxBC ratio. It is possible that rather than being taken up by the tissue, the water insoluble OxBC polymer adhered to the stomach wall, at least partially, resisting ready removal by aqueous washing during processing. Also, apart from stomach, in these tissues and fluids the polymer/apoC ratios in the dosed mice were larger than their counterparts in the control mice.

In contrast, the second group of tissues, comprising liver, large intestine, lung, hamstring, and abdominal muscles, showed no net increase in OxBC levels over those of the controls, despite multiple administrations of a large dose (300 mg/kg body weight). Also, within each tissue, the polymer/apoC ratios were very similar between control and dosed animals. However, whereas liver and large intestine tissues in both dosed and control mice showed polymer/apoC ratios substantially larger than the initial OxBC 4:1 ratio, the ratios in lung, kidney, abdominal muscle, and hamstring muscle were much smaller, at around 1 or even less.

The absence of a net increase in OxBC content and the very similar within-tissue polymer/apoC ratios between control and dosed mice suggest the existence of homeostatic control of polymer and apoC levels in the group 2 tissues. It appears there is either preferential removal of polymer from or enrichment of apoC compounds in lung, kidney, abdominal muscle and especially hamstring. These findings of the existence of significant background OxBC in controls and an apparent limit on accumulation of OxBC further supports the safety of the substance.

There also appears to be selectivity in the retention of individual apocarotenoids in tissues (data not shown). In the apoC fractions, β-ionone was the only apocarotenoid other than DHA that was consistently detected in group 2 tissues. β-ionone was absent from group 1 tissues, except for stomach, especially in dosed mice, and small intestine tissue.

In the OxBC polymer fractions, β-ionone was liberated at detectable levels only in liver and otherwise was undetectable or released only at trace levels in the other tissues and fluids. One other compound was released by NaOH treatment from most of the polymer fractions isolated from the tissues and fluids, except cecum, spleen, and hamstring (data not shown). The compound exhibited the same 153→95 and 153→109 amu MRM transitions as α- and β-cyclocitral but because its HPLC retention time did not correspond to that of either of these compounds its precise identity remains unknown.

The pattern of apocarotenoid release from polymer fractions of tissues and fluids differed from that of OxBC polymer spiked tissues and plasma. Whereas in both situations DHA was the most abundant apocarotenoid released, NaOH treatment of polymer from spiked tissues and plasma also released easily detectable quantities of β-ionone, β-ionone-5,6-epoxide and geronic acid, but no ‘cyclocitral’ (Supplementary Material *S1*), unlike for mouse tissues and fluids. It is not unexpected that some changes may have occurred *in vivo* in the composition of the OxBC polymer during transit from gut to tissues and during any concomitant and subsequent metabolism. Therefore, under these circumstances the values in Table 1 for both polymer and apoC contents are nominal estimates of these fractions that are approximations at best.

The behaviour of the group 1 tissues and fluids reflects processing and elimination of excess dosed OxBC. Urine and intestinal wash contents of dosed mice showed a large excess of OxBC over controls with an accompanying preponderance of polymer, as reflected in the respective high polymer/apoC ratios. In controls, the highest OxBC level in tissues was in liver (67 µg/g) and was largely composed of polymer. Excluding stomach tissue, with the possibility that OxBC was adsorbed to its surface, liver in dosed mice also had the highest OxBC content and a similarly high proportion of polymer. It therefore appears liver preferentially retains OxBC polymer.

In this regard, it is perhaps noteworthy that liver, a major storage site for β-carotene, was reported to have the highest ***concentration*** of β-carotene (50 µg/g) in a high-dose study of uptake and depletion in rats (Shapiro et al., 1984). However, β-carotene uptake for other tissues differed sharply from the OxBC tissue uptake pattern seen in Table 1. In control mice the next highest OxBC contents after liver were in lung (38 µg/g) and hamstring (19 µg/g), compared to the very low β-carotene values seen for lung (0.35 µg/g) and skeletal muscle (0.03 µg/g) in rats in the Shapiro et al., supplementation study.

The control mice results show for the first time that the β-carotene-oxygen copolymer compound and several of the low molecular weight apocarotenoids are naturally present in significant amounts in tissues and fluids. The question arises as to the origin of these compounds.

There are two possibilities. Most likely, OxBC was already present preformed in the mouse diets. It has been established that OxBC occurs naturally in many plant food sources of β-carotene (Burton et al., 2016). Furthermore, it has been shown that the β-carotene-oxygen copolymer compound is the major product formed during loss of β-carotene in a large variety of plant food items (Schaub et al., 2017). Given that the spontaneous formation in air of the β-carotene polymer occurs regardless of the environment, plant or chemical, and apocarotenoids are inevitably formed as a side product (Burton et al., 2014; Mogg and Burton, 2021), the similar polymer/apoC ratio of ∼4 seen for both control and dosed stomach tissue is consistent with a dietary source of OxBC.

The second possibility is that dietary β-carotene is subject to non-enzymatic oxidation *in vivo*. It is known that mice and rats efficiently convert β-carotene to vitamin A in the gut but absorb carotenoids intact only when they are provided in the diet at high, non-physiological levels (Goodman et al., 1966; Huang and Goodman, 1965; Lee et al., 1999; Shapiro et al., 1984). However, the level of dietary β-carotene in the mouse diets was quite low. For example, the Charles River Rat and Mouse diet contains 0.8 ppm β-carotene. Furthermore, most of the dietary β-carotene would be converted into vitamin A (Goodman et al., 1966; Huang and Goodman, 1965).

## 4. Conclusions

The toxicology study in rats established an MTD of 5,000 mg/kg, an LD_50_ of 30,079 mg/kg and a NOAEL of 1875 mg/kg body weight. Synthetic OxBC has been used for livestock, pets, and humans at a level of approximately 0.5 mg/kg body weight/day, which is 3,750-fold below the NOAEL-based threshold. The absence of evidence of toxicity with chronic dosing at the NOAEL level in the toxicology study indicates that oral use of synthetic OxBC at levels of ∼0.5 mg/kg body weight has a wide safety margin.

The OxBC uptake study established that OxBC in both its polymer and apoC forms was naturally present in all mouse tissues and fluids examined. The highest background levels were in liver, lung, and hamstring. Supplemental OxBC, even at the high dose level used in the study, barely affected the net background levels in liver, kidney, lung, and muscle tissues after 2- and 5-days daily dosing. This finding suggests there is control of tissue OxBC levels. Furthermore, whereas the polymer fraction was enriched relative to the apoC fraction in liver, implying preferential retention, it was depleted in lung, kidney, hamstring, and abdominal muscle.

However, net increases in OxBC and the polymer fraction did occur in the 2-5-day dosing period in urine, intestinal content, plasma, feces, spleen, and cecum, which is consistent with processing of OxBC and preferential on-going elimination of the polymer at these sites.

The discovery of the background presence of OxBC in all examined mouse tissues and fluids at levels that were barely affected by dosing in liver, lung, kidney, and muscle suggests the existence of a safety mechanism for limiting OxBC exposure in at least several major tissues.

## Abbreviations

OxBC: fully oxidized β-carotene
apoC: apocarotenoid
DHA: dihydroactinidiolide
MTD: maximum tolerated dose
MRM: multiple reaction monitoring
NOAEL: no observed adverse effect level
NRC: National Research Council of Canada
PEG: polyethylene glycol

## Acknowledgment

Mr. Daniel Trudeau and NRC IRAP are thanked for their invaluable support.

