## Supplemental File S1 for "Safety and uptake of fully oxidized β-carotene"

### Development of an LC-MS method for determination of OxBC polymeric and apocarotenoid fractions

*Abbreviations*: APCI: Atmospheric pressure chemical ionization; CE: collision energy; ESI: electrospray ionization; GPC: gel phase chromatography; HRMS: high resolution mass spectrometry; MALDI: matrix-assisted laser desorption/ionization; MRM: multiple reaction monitoring; OxBC: fully oxidized β-carotene; OxBC apoC: OxBC apocarotenoid fraction; TIC: total ion chromatogram.

#### 1. Objective

To develop a mass spectrometric method to analyze OxBC's two principal components, the major polymeric fraction (OxBC polymer) and the minor monomeric apocarotenoid fraction (OxBC apoC), for application to the semi-quantitative determination of the levels of these components in animal tissues and fluids.

#### 2. Methods and materials

The OxBC polymer and OxBC apoC fractions were isolated from OxBC, prepared as reported earlier (Burton et al., 2014), and as described further in section 2.5.1 of this parent paper. OxBC polymer was precipitated by dropwise addition of hexane to an ethyl acetate solution of OxBC, repeating the process four times. Solutions of OxBC apoC and OxBC polymer were prepared in methanol (10 mg/mL) for analysis by LC-MS and GPC-MS.

Mouse serum, liver, plasma, and muscle tissues were obtained from BioIVT (Hicksville, NY). All materials were pooled gender homogenates from non-medicated and non-immunized Balb/C animals. Serum, plasma, liver and muscle tissues were stored at -70°C. When used, samples were thawed at room temperature and kept on ice in the fridge for no more than 48 h.

##### 2.1. LC-MS, GPC-MS, and LC-MRM analysis

LC-MS analysis of OxBC apoC was carried out using a reverse phase Atlantis T3 column (1 x 150 mm, 3 μm) and a binary gradient program with buffer A: 0.1 % formic acid in water and buffer B: 0.1% formic acid in acetonitrile. The LC gradient used was: 0 to 3 min, B = 20%; 3 to 25 min, linear gradient to B = 95%; 25 to 32 min, gradient held at B = 95%; 32.1 to 40 min, equilibration at B = 20%. The flow rate was 50 μL/min.

GPC-MS analysis of the OxBC polymer was performed on a Jordi X-Stream column (1 mm x 25 cm) with 0.1 % formic acid in THF mobile phase and a flow rate of 40 μL/min

LC-MRM was performed using a 4000 QTrap instrument (AB Sciex, Concord, ON) coupled to an Ultimate 3000 UHPLC (Waltham, MA). High-resolution MS was performed using a Thermo LTQ Orbitrap XL-ETD instrument (Waltham, MA).

All reagents and solvents used were obtained from Sigma-Aldrich (Oakville, ON) and were of the highest available purity.

The method development process was carried out in several stages, analyzing the OxBC apoC fraction first and then the OxBC polymer fraction.

##### 2.2. OxBC apoC fraction

LC-MS profiling of the apocarotenoids in the OxBC apoC fraction was carried out with a low-resolution full scan of on-column-injected OxBC apoC on the Sciex 4000 Qtrap and high-resolution mass spectrometry (Orbitrap) to confirm the chemical formulas of detected ions.

A sample of OxBC apoC was analyzed by LC-MRM to identify individual apocarotenoids and to provide a reference for serum spiked with OxBC apoC. Serum (1 mL) was spiked with OxBC apo in methanol (100 µL, 10 mg/mL), vortexed for several minutes, dried by vacuum centrifugation at 35°C for several hours, then extracted by shaking with methanol (100 µL). After centrifugation, a sample of the supernatant (~100 µL) was transferred to a glass vial for LC-MRM analysis.

##### 2.3. OxBC polymer fraction

The intact OxBC polymer was analyzed using MALDI-MS and GPC-UV-MS using both electrospray ionization (ESI-MS) and atmospheric pressure chemical ionization (APCI-MS).

***Apocarotenoid marker compounds released by chemical treatment of the OxBC polymer***

Because treatment of OxBC polymer with sodium hydroxide had already been shown to release various apocarotenoid compounds (Mogg and Burton, 2021), use was made of this knowledge to apply the LC-MS method to NaOH-treated OxBC polymer to identify liberated apocarotenoids that could serve as marker compounds for the polymer.

Given that it was known the original isolated OxBC polymer starting compound could still contain traces of apocarotenoid compounds, it was important to remove these residual low molecular weight contaminants. As both the OxBC apoC and OxBC polymer fractions are soluble in ethanol but only the apoC apocarotenoid components are soluble in hexane, the apoC compounds could be removed by a method using hexane to precipitate the polymer from a concentrated solution of the compound in ethyl acetate, leaving the apocarotenoids in solution. Hexane precipitations were carried out for a total of six times.

Chemical treatment of the purified polymer was carried out by adding aqueous NaOH (1 M, 240 µL) to a solution of the polymer in methanol (1 mL) and heating at 75°C for 4 hrs with shaking. After cooling to room temperature, the sample was acidified dropwise with HCl (2 M) to approximately pH 4 and transferred directly to a glass vial for LC-MRM analysis.

###### 2.3.1. LC-MS analysis of mouse tissues and plasma spiked with OxBC polymer

The LC-MS method was used to evaluate the ability to recover apocarotenoids from mouse tissue homogenates and plasma spiked with OxBC polymer. The process was as follows:

1. OxBC polymer (20 mg) dissolved in methanol (1 mL) was added to muscle or liver homogenate (1 mL; 0.2 mg tissue/mL) or plasma (1 mL) and vortex-mixed with ethanol (1 mL) for 1 min. Hexane (1 mL) was then added and the mixture vortex-mixed for 1 min. The phases were separated by centrifugation and the hexane layer removed. Hexane (1 mL) was added to the remaining aqueous alcohol layer and the procedure repeated. The

recovered hexane layers were combined, evaporated to dryness and the residue dissolved in methanol (1 mL) for LC-MS analysis (Hexane Wash 1).

2. Ethyl acetate (1 mL) and water (1 mL) were added to the remaining aqueous alcohol fraction and vortex mixed for 1 min. After the ethyl acetate layer was removed, the remaining aqueous fraction was extracted in the same manner with ethyl acetate (1 mL). The ethyl acetate fractions were combined and dried down to give the OxBC polymer pellet.
3. The recovered pellet was redissolved in ethyl acetate (50 µL) using sonication and vortex mixing. Hexane (1 mL) was added slowly to precipitate the OxBC polymer. After standing for 10 min the precipitate was spun down and the hexane layer decanted. The recovered precipitate was subjected to the hexane washing and precipitation procedure five times. The combined hexane fractions were evaporated to dryness and the residue dissolved in methanol (1 mL) for LC-MS analysis (Hexane Wash 2).
4. The hexane washed OxBC polymer pellet was dissolved in methanol (1 mL). A sample (100 µL) was used for LC-MS analysis. The remaining solution (900 µL) was treated with aqueous sodium hydroxide (1 M, 240 µL) by heating at 75°C for 4 hrs with shaking and the total volume (1140 µL) analyzed directly by LC-MS.

OxBC polymer also was incubated with plasma for 3 h at room temperature to determine the polymer's stability towards breakdown, for example by plasma enzymes.

#### 3. Results and Discussion

##### 3.1. OxBC apoC fraction

Figure 1 shows a typical LC-MS chromatogram for on-column injected OxBC apoC analyzed by low resolution MS using the 4000 QTrap triple-quadrupole mass spectrometer.

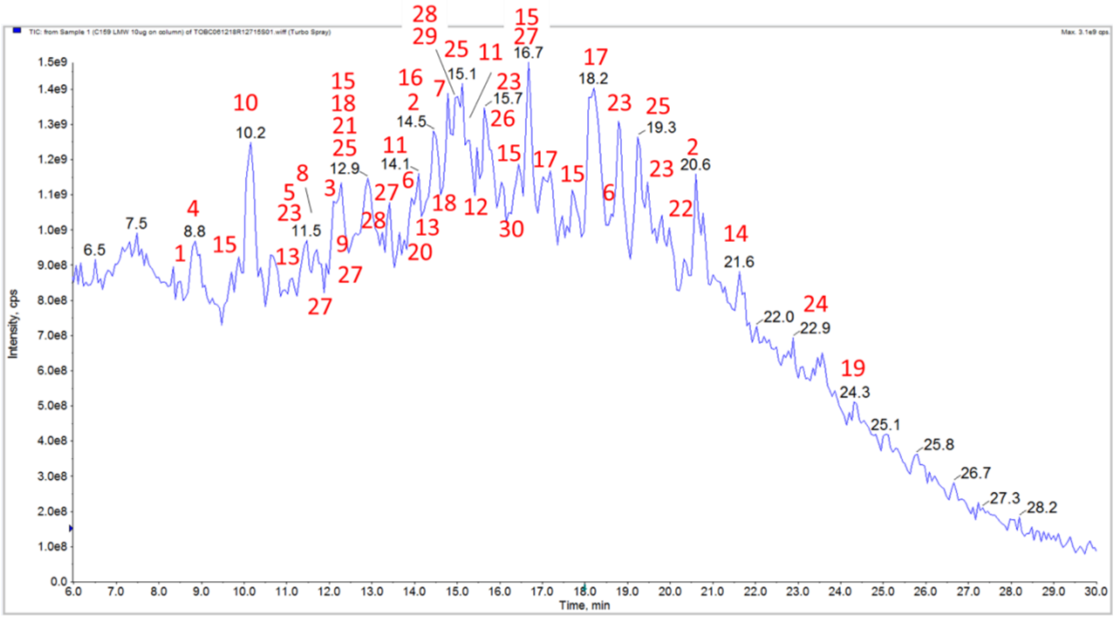


**Figure 1.** LC-MS profile with a low-resolution full scan of on-column OxBC apoC (10 µg) on the Sciex 4000 Qtrap. The numbers in red refer to the compounds listed in Table 1 and their chemical structures in the Appendix. Numbers in black are retention times.

To confirm the chemical formulas of detected ions, OxBC apoC also was analysed using the Orbitrap high-resolution mass spectrometer (HRMS). The apocarotenoid compounds identified are shown in Table 1. Elemental compositions were obtained using the Orbitrap's built-in algorithm (Thermo Xcalibur Qual Browser, Thermo Corp.) with a cut-off mass accuracy of 15 parts-per-million (ppm). Also shown are the nominal masses observed using the 4000 QTrap low resolution triple-quadruple instrument's observed adduct ions ($[M+H]^+$, $[M+NH_4]^+$), as well as the two fragment ions (Q3-1, Q3-2) used for further MRM experiments.

**Table 1.** Selected apocarotenoid compounds identified in OxBC apoC by LC-MS with both low-resolution MS (QTrap) and high-resolution mass spectrometry (Orbitrap). The two transitions monitored for each compound by MRM are shown as Q3-1 and Q3-2. CE = collision energy. Structures of compounds 1-32 are given in the Appendix.

| | | | Qtrap | | | | OrbiTrap HRMS | | | Optimized Multiple Reaction Monitoring (MRM) Parameters | | | | |
|---|---|---|---|---|---|---|---|---|---|---|---|---|---|---|
| Peak | Compound Number | Name | $[M+H]^+$ | $[M+NH_4]^+$ | RT (min.) | Formula $[M+H]^+$ | Calc. MW [M+H]+ | Obs. MW $[M+H]^+$ | Mass Error ppm | Q1 | Q3-1 | CE | Q3-2 | CE |
| | (See Appendix) | | | * | | | | | | | | | | |
| 1 | **11** | 2-Methyl-6-oxo-2,4-heptadienal | 139.1 | 156.1 | 8.4 | $C_8H_{10}O_2$ | 139.0759 | 139.0754 | -3.6 | -- | -- | -- | -- | -- |
| 3 | **2** | ß-Cyclocitral | 153.1 | 170.1 | 12.1 | $C_{10}H_{17}O_1$ | 153.1279 | 153.1283 | 2.6 | 153.1 | 95.1 | 20 | 109.1 | 20 |
| 5 | **27** | 2-Hydroxy-2,6,6-trimethylcyclohexanone | 157.1 | 174.1 | 11.4 | $C_9H_{16}O_2$ | 157.1228 | 157.1224 | -2.5 | 157.1 | 69.1 | 20 | 111.2 | 15 |
| 7 | **24** | 4-Oxo-ß-cyclocitral | 167.1 | 184.1 | 15.0 | $C_{10}H_{15}O_2$ | 167.1072 | 167.1068 | -2.4 | 167.1 | 121.1 | 25 | 149.1 | 22 |
| 8 | **25** | 2,6,6-Trimethyl-1-cyclohexene-1-carboxylic acid | 169.1 | 186.1 | 11.7 | $C_{10}H_{17}O_2$ | 169.1228 | 169.1226 | -1.2 | 169.1 | 80.8 | 30 | 123.1 | 20 |
| 10 | **32** | Geronic acid | 173.2 | 190.2 | 10.0 | $C_9H_{17}O_3$ | 173.1178 | 173.11714 | -3.8 | 173.2 | 109.1 | 20 | 127 | 15 |
| 12 | **8** | Dihydroactinidiolide | 181.1 | 198.1 | 15.3 | $C_{11}H_{17}O_2$ | 181.1228 | 181.1223 | -2.8 | 181.1 | 107.1 | 30 | 135.1 | 30 |
| 14 | **4** | ß-ionone | 193.1 | 210.1 | 21.6 | $C_{13}H_{21}O$ | 193.1592 | 193.1587 | -2.6 | 193.1 | 109.1 | 25 | 135.1 | 20 |
| 16 | **6** | 4-Oxo-ß-ionone | 207.1 | 224.1 | 14.5 | $C_{13}H_{19}O_2$ | 207.1385 | 207.1381 | -1.9 | 207.1 | 111 | 20 | 165.3 | 17 |
| 17 | **5** | ß-Ionone-5,6-epoxide | 209.1 | 226.1 | 18.2 | $C_{13}H_{21}O_2$ | 209.1542 | 209.1539 | -1.4 | 209.1 | 85 | 20 | 151.1 | 14 |
| 19 | **22** | ß-Ionylidene acetaldehyde | 219.1 | 236.1 | 24.3 | $C_{15}H_{23}O_1$ | 219.1749 | 219.1744 | -2.3 | 219.1 | 163.2 | 20 | 191.1 | 15 |
| 20 | **12** | 6,6-Dimethylundec-3-en-2,5,10-trione | 225.1 | 242.1 | 14.0 | $C_{13}H_{21}O_3$ | 225.1491 | 225.1487 | -1.8 | 225.1 | 107.1 | 25 | 165.3 | 15 |
| 22 | **18** | ß-Ionylidene acetaldehyde-5,6-epoxide | 235.1 | 252.1 | 20.6 | $C_{15}H_{23}O_2$ | 235.1698 | 235.1693 | -2.1 | 235.1 | 139.3 | 20 | 165.1 | 22 |
| 24 | **20** | ß-apo-13-Carotenone-5,6-epoxide | 275.1 | 292.1 | 23.1 | $C_{18}H_{27}O_2$ | 275.2011 | 275.2008 | -1.1 | 275.1 | 181.3 | 19 | 191.2 | 25 |

*Some peaks also show adducts with acetonitrile $[M + ACN]^+$ and/or methanol $[M + MeOH]^+$; Data not listed

MRM is the accepted standard for small molecule quantitation in complex mixtures. To optimize MRM parameters, product ion scans were first performed using arbitrary collision energies (CEs) and unique fragment ions were chosen for each compound. CE optimization was then performed by ramping the CE in small increments: Optimum CEs were chosen based on the maximum signal intensity observed for each MRM transition monitored.

##### 3.1.1. LC-MRM analysis of OxBC apoC standard and spiked serum

Fig. 2 shows that the LC-MRM method applied to the OxBC apoC standard yielded a clean, greatly simplified chromatogram with three main apocarotenoid peaks: dihydroactinidiolide

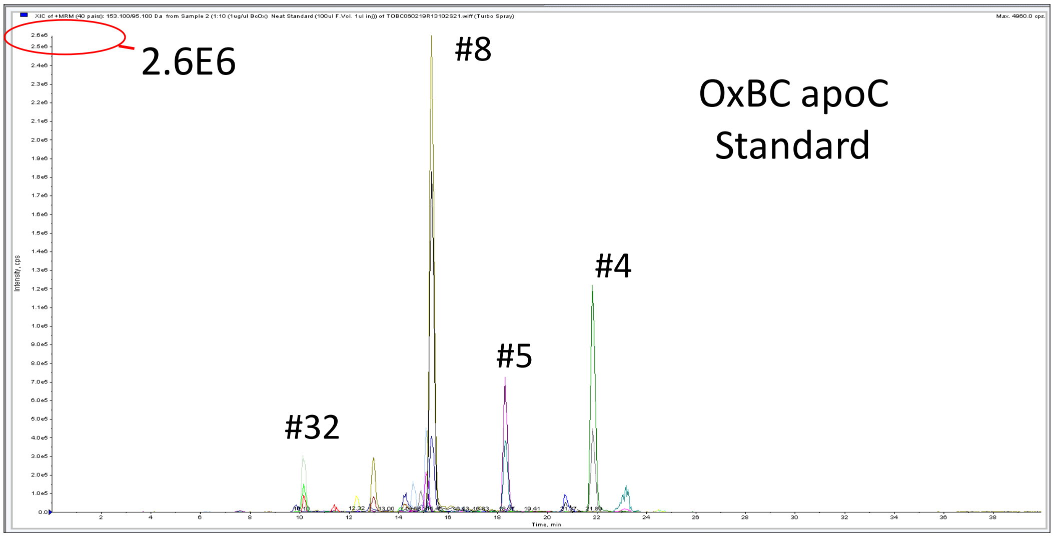


**Figure 2.** LC-MRM analysis of OxBC apoC standard. The three biggest peaks in order of elution are dihydroactinidiolide (DHA, compound #8; 15.3 min.), ß-ionone-5,6-epoxide (compound #5; 18.2 min.) and ß-ionone (compound #4; 21.6 min.). Compound #32 is geronic acid.

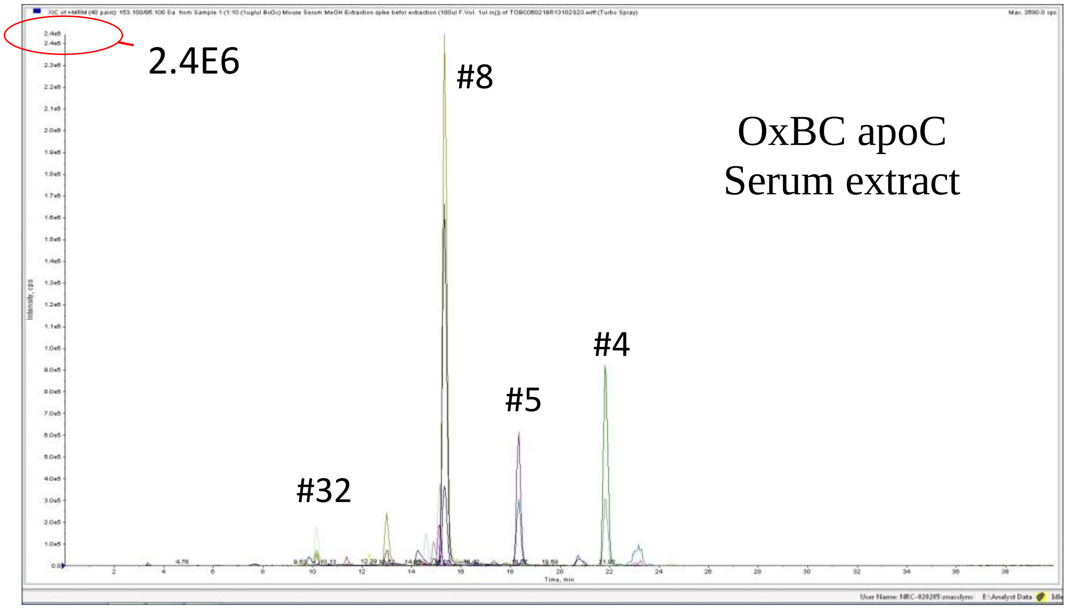


**Figure 3**. LC-MRM analysis of methanol extract of vacuum-dried OxBC apoC spiked serum. The three major peaks in order of elution are dihydroactinidiolide (compound #8), ß-ionone-5,6-epoxide (compound #5) and ß-ionone (compound #4).
(DHA, compound #8), ß-ionone-5,6-epoxide (compound #5) and ß-ionone (compound #4). Compound #32 is geronic acid (chemical structures are provided in the Appendix)

Fig. 3 shows that the MRM chromatogram of the apocarotenoids recovered by methanol extraction of vacuum dried OxBC apoC spiked serum is very similar to that of the OxBC apoC standard itself. Recoveries from serum of individual carotenoids were performed in triplicate at three concentration levels of 10, 100 and 1000 ng. Recoveries ranged from 50% to 130%, as calculated from peak intensities.

##### 3.2. OxBC polymer fraction

Attempts to use MALDI-MS to observe the intact OxBC polymer were unsuccessful (data not shown). Therefore, a GPC-UV-MS approach was adopted to attempt to monitor the intact polymer using both electrospray ionization (ESI-MS) and atmospheric pressure chemical ionization (APCI-MS).

The GPC-UV-MS results obtained for OxBC polymer using ESI and APCI modes are shown in Figs. 4 and 5, respectively. No meaningful data were obtained, which might possibly have shown multiply charged clusters of ions corresponding to the presence of polymers, if ionisation behaviour resembled that of a small protein.

Detection of the polymer through in-source fragmentation was also attempted, wherein a high voltage was applied at the entrance of the mass spectrometer to generate fragments from larger polymers, which could then be characterised (Figure 5C). This approach did not generate any small repeat units as typically would be observed for a polysaccharide.

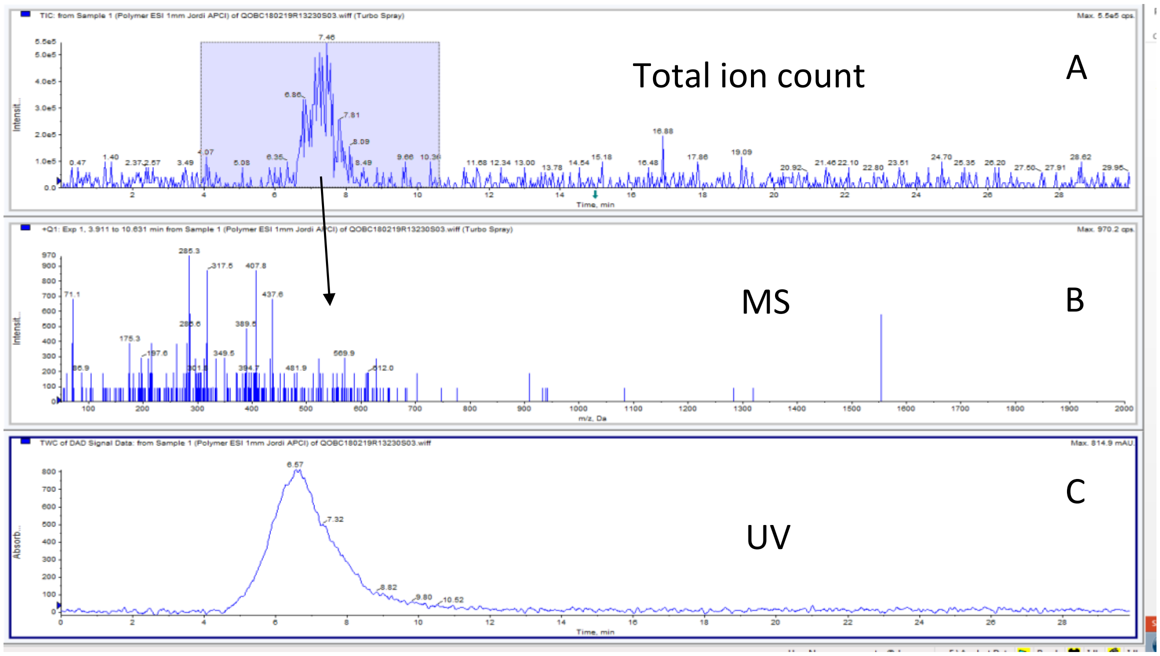


**Figure 4.** GPC-UV-MS analysis of the OxBC polymer using ESI. (A) Total ion count (TIC); (B) MS spectrum extracted from the time window highlighted in A; (C) GPC-UV trace.

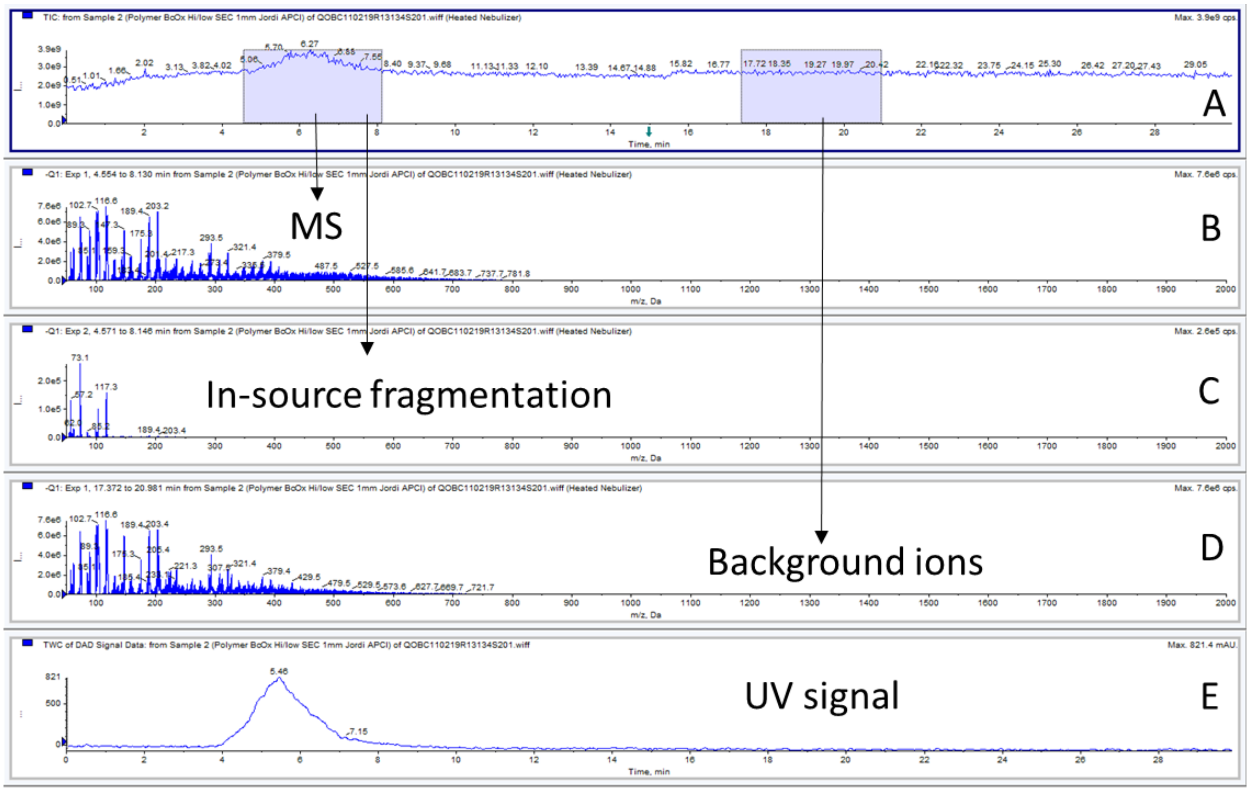


**Figure 5**. GPC-UV-MS analysis of OxBC polymer using APCI. (A) TIC; (B) Extracted MS spectrum from the highlighted time window, showing no multiply charged ions; (C) Extracted spectrum of in-source fragmentation MS scan of the OxBC polymer gave no information on any polymer repeat units; (D) Extracted MS spectrum confirming that most singly charged ions observed in B originate from background; (E) GPC-UV trace.

Given the polymer's unique, complex chemical composition, with many chemical subunits, it is not unexpected that mass spectrometric detection was difficult.

In summary, GPC-MS analysis using APCI or ESI interfaces did not detect intact OxBC polymer.

##### 3.2.1. Indirect LC-MRM detection of OxBC polymer using sodium hydroxide treatment

An LC-MRM method was successfully developed for indirect detection of the OxBC polymer using analysis of apocarotenoids released by treatment with sodium hydroxide.

The LC-MS (HRMS) of OxBC polymer treated with sodium hydroxide for 4 h at 75°C is shown in Fig. 6. Analysis with the 4000 QTrap low-resolution mass spectrometer was used to develop a LC-MRM-based quantitative assay. As anticipated from previous work (Mogg and Burton, 2021), NaOH degradation gave rise to apocarotenoids that are also seen in the OxBC apoC fraction. Selected identified products are listed in Table 2. No products unique to the polymer were detected.

##### 3.2.2. Detection of OxBC polymer in spiked tissue homogenates and plasma

The hexane-ethyl acetate-water extraction and precipitation procedure, together with NaOH treatment, in combination with the LC-MRM method revealed that OxBC polymer could be identified by detecting apocarotenoids in plasma and homogenates of muscle and liver spiked with the OxBC polymer fraction.

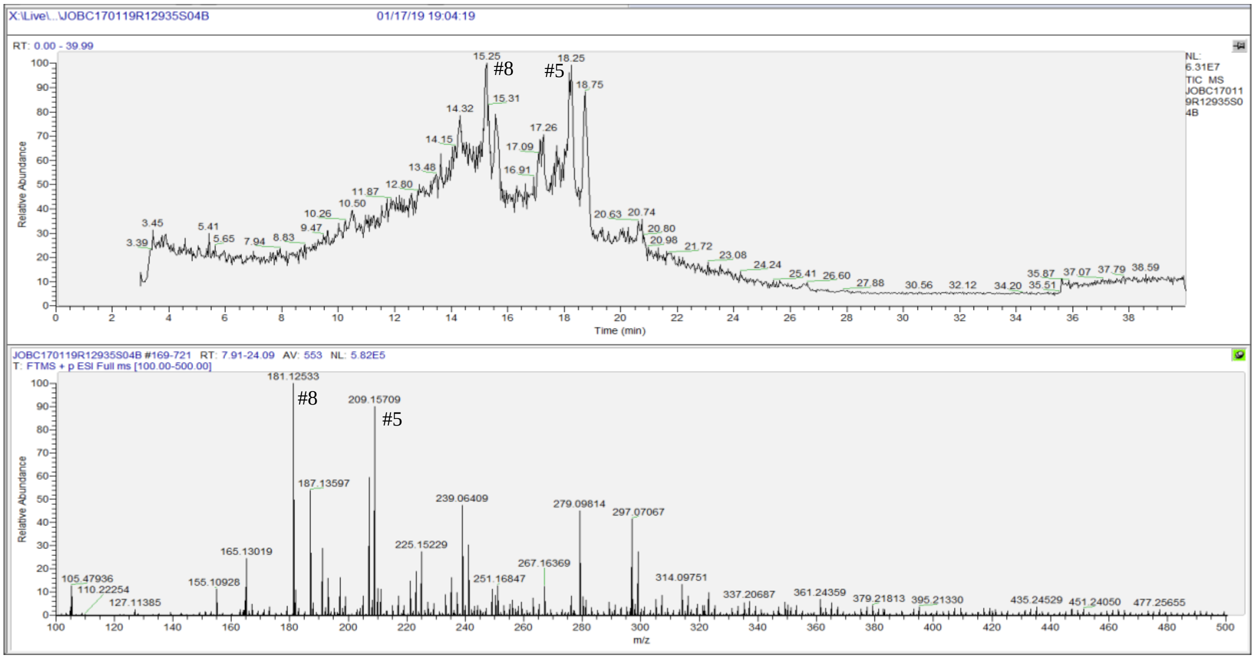


**Figure 6**. LC-MS (HRMS) trace for NaOH degraded OxBC polymer and extracted MS spectrum between 7.9 min and 24 min. #8: dihydroactinidiolide (DHA), #5: ß-ionone-5,6-epoxide

**Table 2.** Selected compounds identified in NaOH-degraded OxBC polymer.

| Qtrap | | Orbitrap HRMS validation | | | Mass Error | Compound[2] |
|---|---|---|---|---|---|---|
| $[M+H]^+$ | Retention Time[1] | Formula | Calc. MW $[M+H]^+$ | Obs. MW $[M+H]^+$ | ppm | |
| 181.1 | 15.2 | $C_{11}H_{17}O_2$ | 181.1229 | 181.1253 | 13.4 | #8 |
| 193.2 | 21.7 | $C_{13}H_{21}O$ | 193.1592 | 193.1619 | 13.9 | #4 |
| 207.1 | 14.5 | $C_{13}H_{19}O_2$ | 207.1385 | 207.1414 | 14.0 | #6 |
| 209.1 | 18.3 | $C_{13}H_{21}O_2$ | 209.1542 | 209.1571 | 13.9 | #5 |

[1] 40-minute runs

[2] #4: ß-ionone; #5: ß-ionone-5,6-epoxide; #6: 4-oxo-ß-ionone; #8: dihydroactinidiolide (DHA). See the Appendix for chemical structures.

##### 3.2.3. Detection of OxBC polymer in spiked muscle and liver homogenates

OxBC polymer purified by precipitating OxBC from ethyl acetate solution with hexane six times still contained traces of apocarotenoid compounds, as seen for the OxBC polymer standard in Fig 7A. Extensive washing of the polymer pellet recovered from spiked tissue homogenate using successive ethyl acetate/hexane precipitations removed much of the contaminating low molecular weight hexane-soluble compounds, as shown for muscle homogenate in Fig. 7B and

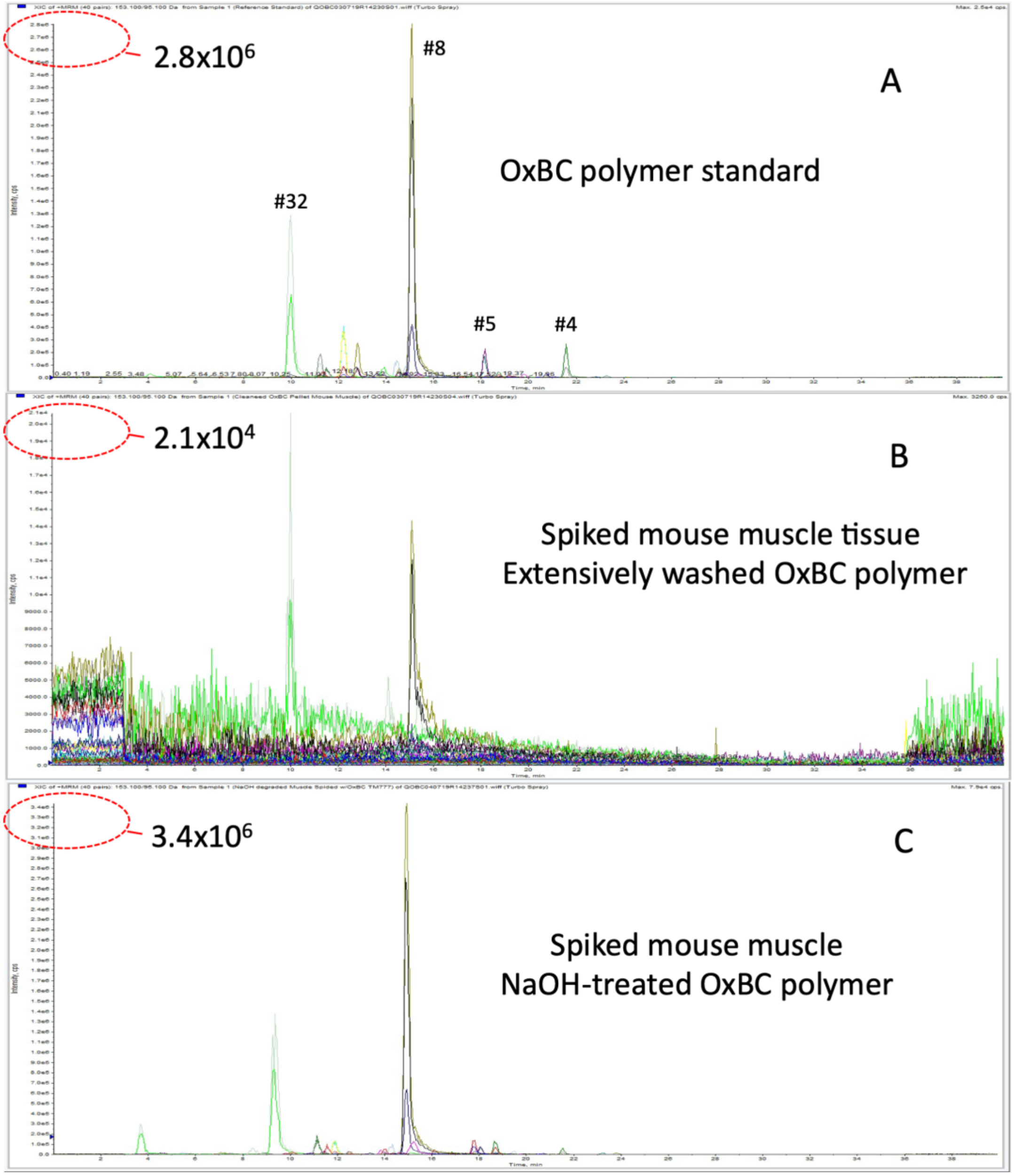


**Figure 7.** Results for spiking of mouse muscle tissues with OxBC polymer showing: (A) the presence of residual apocarotenoids in the intact polymer standard; (B) apocarotenoids remaining after extensive washing of the precipitated polymer with hexane, and (C) apocarotenoids released after NaOH treatment of the hexane washed OxBC polymer in (B). Peak numbers refer to structures listed in the Appendix.

the presence of apocarotenoids in Hexane Washes 1 and 2 (Fig. 8). The validity of this method depends upon the apocarotenoids and polymer both being soluble in ethyl acetate but only the apocarotenoids being soluble in hexane.

Liver homogenate spiked with OxBC polymer also was examined. The results for extraction of OxBC polymer as determined by apocarotenoid detection was essentially the same as for muscle homogenate.

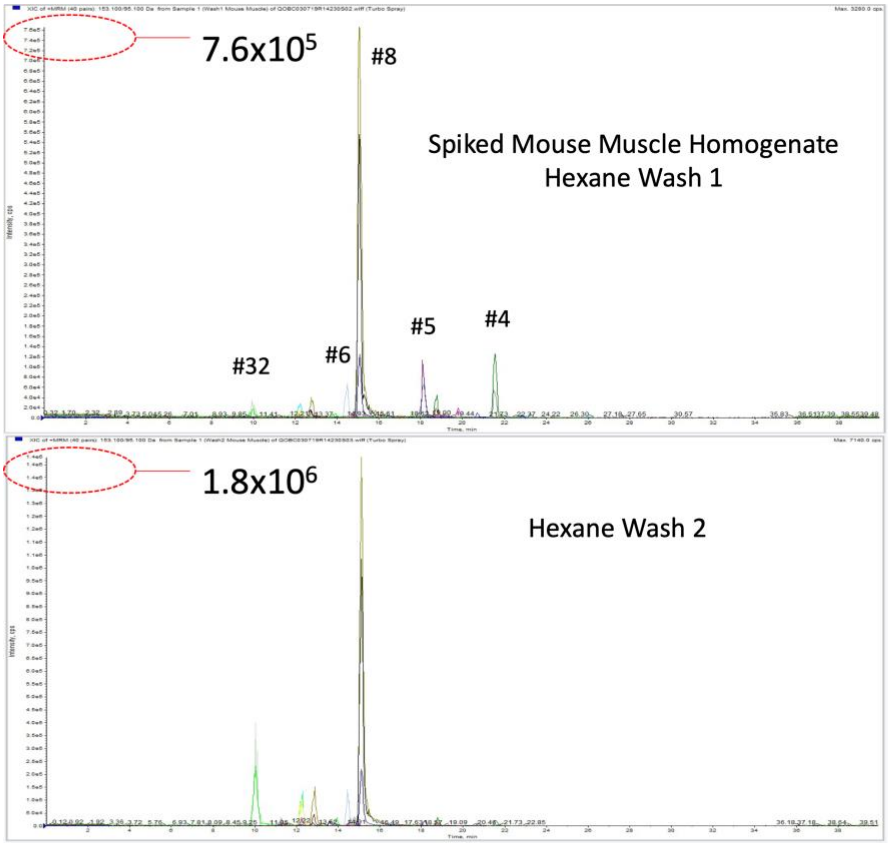


**Figure 8**. Apocarotenoids extracted into combined hexane extracts of spiked mouse muscle homogenate (Hexane Wash 1) and in combined hexane extracts after extensive washing of the precipitated OxBC polymer pellet (Hexane Wash 2). (See Section 2.3.1.)

##### 3.2.4. Detection of OxBC polymer in spiked plasma

Mouse plasma was spiked with OxBC polymer and processed in a manner very similar to that used for detecting OxBC polymer in spiked muscle and liver tissue homogenates. In addition, the stability of OxBC polymer in plasma was assessed by comparing extractions immediately after spiking (0 h) and after standing at room temperature for 3 h.

Fig. 9 shows the results for extraction at 0 h. There was a significant decrease in signal intensity and signal-to-noise ratio of all apocarotenoid peaks in the OxBC polymer recovered from spiked plasma after extensive washing with hexane (Fig. 9B) compared to the OxBC polymer standard (Fig. 9A. However, after NaOH treatment of the OxBC polymer the signal intensities of the apocarotenoids were almost four times the levels observed in the standard (Figs. 9C vs. 9A) and more than 10-fold greater than those in the washed pellet (Figs. 9C vs. 9B). It is concluded therefore that the increased levels of apocarotenoids in the NaOH-treated polymer sample originate from NaOH degradation of the polymer given that the polymer itself is not directly detectable.

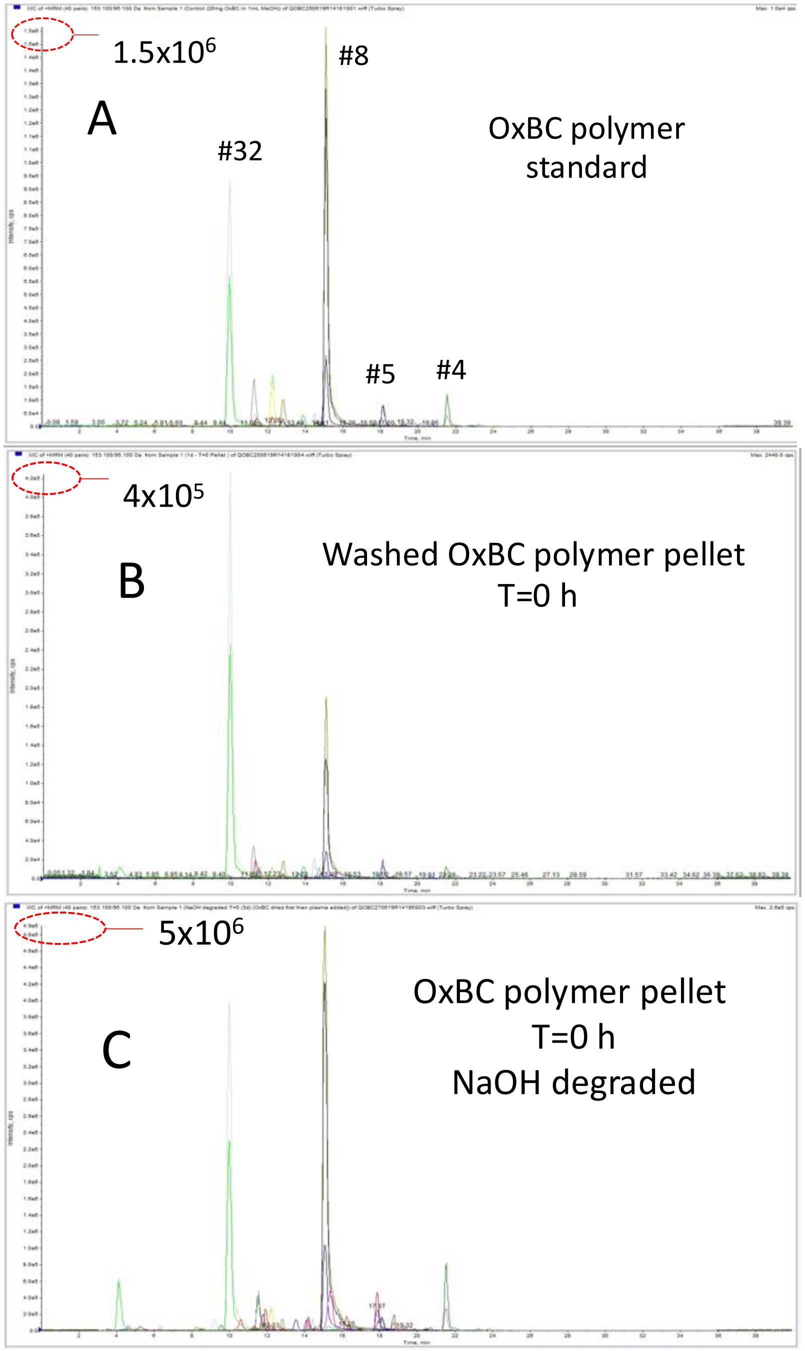


**Figure 9.** Mouse plasma spiked with OxBC polymer, vortex mixed and immediately extracted with hexane (T=0 h). (A) Apocarotenoid compounds detected in the intact OxBC polymer standard; (B) apocarotenoid compounds remaining after extensive extraction with hexane; (C) apocarotenoid compounds released after NaOH treatment of hexane washed OxBC polymer. Peak numbers refer to structures in the Appendix. **Note:** See Fig. 10 for chromatograms of Hexane Wash solutions containing extracted apocarotenoids.

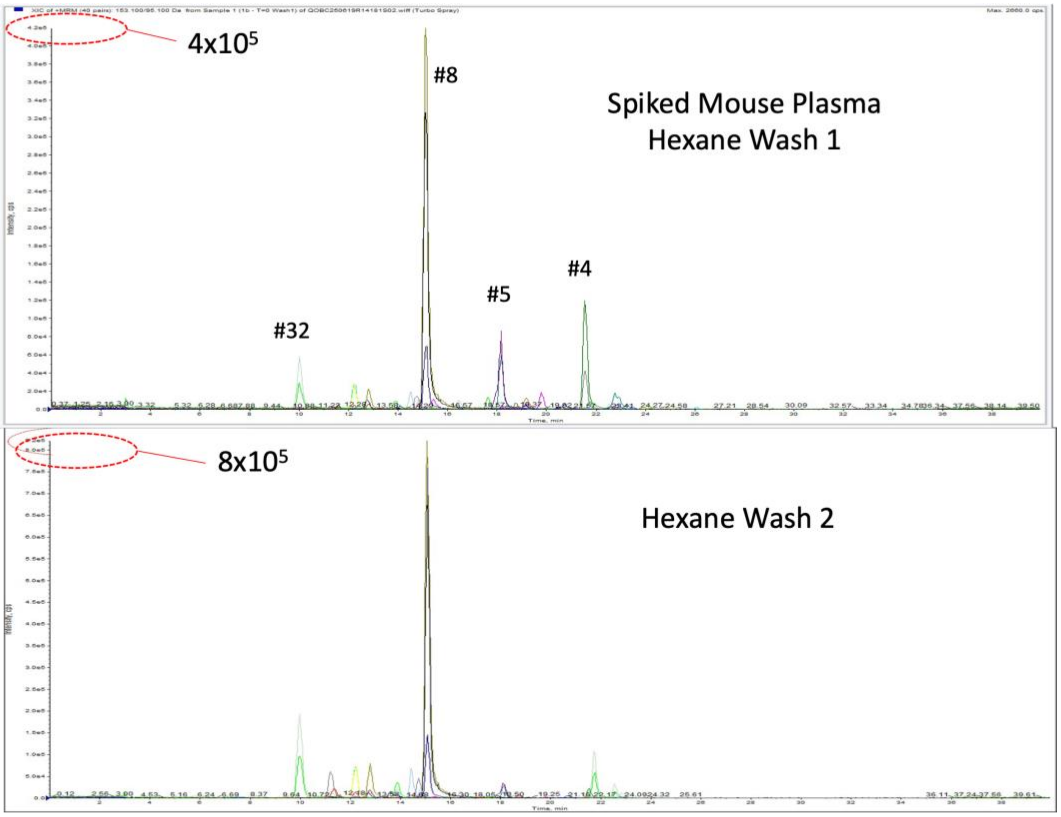


**Figure 10**. Apocarotenoids initially extracted into hexane from spiked mouse plasma (Hexane Wash 1) and apocarotenoids extracted by extensive hexane washing of isolated polymer pellet (Hexane Wash 2).

There was no evidence of any changes in the LC-MRM chromatograms after incubation of OxBC polymer with plasma for 3 hours and processing of the sample in the same manner as for the T=0 h sample. This result indicated that the polymer was stable under the conditions and not readily susceptible to degradation, for example, by plasma enzymes.

Strong evidence supporting the validity of the process using the NaOH treatment to detect polymer was obtained using a second method to spike plasma with OxBC polymer. Instead of mixing plasma with a solution of OxBC polymer in methanol, the OxBC polymer solution was first dried by evaporation of the methanol before thoroughly incubating the dry residue with plasma. Extraction with hexane of the plasma spiked in this manner and extensive washing of the ethanol-precipitated OxBC polymer pellet in the usual manner, followed by LC-MRM analysis of the washed polymer pellet, indicated that the apocarotenoids seen previously were not present at detectable levels (Fig. 11B). However, LC-MRM analysis of the hexane washed pellet subsequently treated with NaOH clearly showed the presence of NaOH-released apocarotenoids (Fig. 11C). It appears the OxBC polymer and the trace residual apocarotenoids it contained were fully taken up into the intact lipoprotein structures and that the original contaminating free apocarotenoids were stripped from the polymer by being fully extracted into hexane after the plasma was denatured with ethanol. The same result was obtained when the dried OxBC polymer was incubated with plasma for 3 h at room temperature, confirming the stability of the polymer under these conditions during this period.

LC-MRM analysis of un-spiked plasma samples confirmed the absence of any interfering background ions.

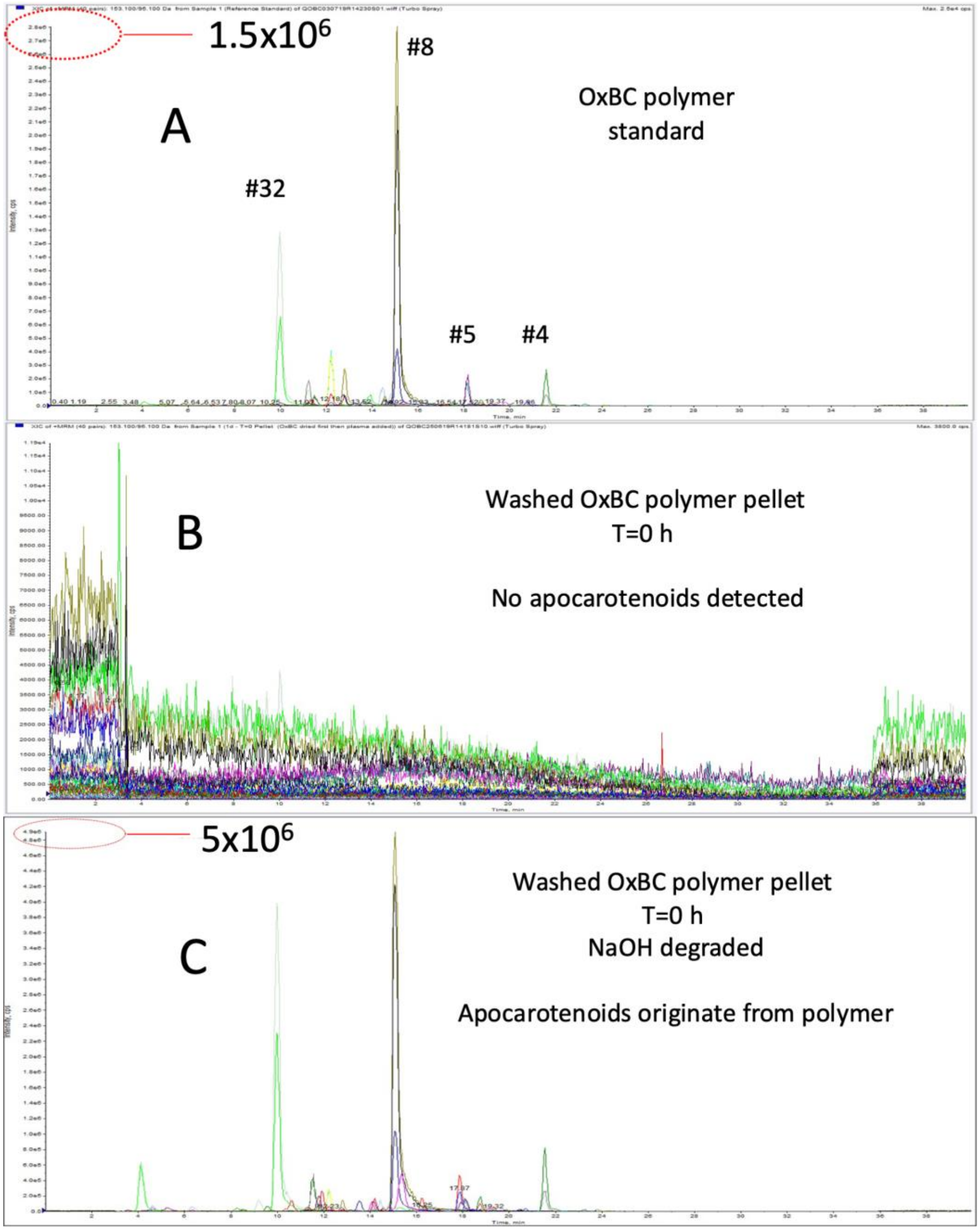


**Figure 11.** Mouse plasma spiked using dried OxBC polymer at T=0 h (approach 2), showing: (A) apocarotenoids observed in the intact standard; (B) absence of apocarotenoids after hexane extraction of plasma; (C) apocarotenoids released by NaOH degradation of isolated OxBC polymer.

#### 4. Conclusion

LC-MS analysis of the OxBC apoC fraction confirmed the presence of a multitude of apocarotenoids. LC-MRM analysis yielded a greatly simplified chromatogram containing four of the most abundant OxBC apocarotenoid compounds identified previously (Burton et al., 2014), namely DHA, ß-ionone, ß-ionone-5,6-epoxide and geronic acid. The same pattern of detected compounds was obtained for plasma spiked with OxBC apoC.

The OxBC polymer was not amenable to direct analysis by mass spectrometry. However, NaOH treatment of the polymer generated the same four apocarotenoids seen in OxBC apoC, as detected by LC-MRM analysis. Successful indirect detection of OxBC polymer in this manner in spiked muscle and liver homogenates and plasma confirmed NaOH treatment afforded a viable approach to the detection and potential quantifying of OxBC polymer in tissues and body fluids.

**Appendix**. Partial list of structures assigned to compounds in the OxBC apoC fraction.

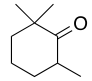
**1**
2,2,6-trimethylcyclohexanone
$C_9H_{16}O$ 140.12

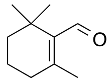
**2**
β-Cyclocitral
$C_{10}H_{16}O$ 152.12

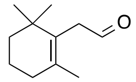
**3**
β-Homocyclocitral
$C_{11}H_{18}O$ 166.14

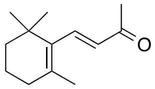
**4**
β-Ionone
$C_{13}H_{20}O$ 192.15

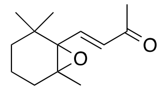
**5**
β-Ionone-5,6-epoxide
$C_{13}H_{20}O_2$ 208.15

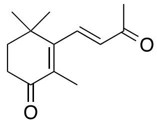
**6**
4-Oxo-β-ionone
$C_{13}H_{18}O_2$ 206.13

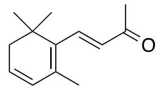
**7**
4-(2,6,6-Trimethylcyclohexa-1,3-dienyl)but-3-en-2-one
$C_{13}H_{18}O$ 190.14

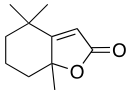
**8**
Dihydroactinidiolide
$C_{11}H_{16}O_2$ 180.12

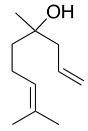
**9**
4,8-Dimethyl-1,7-nonadiene-4-ol
$C_{11}H_{20}O$ 168.15

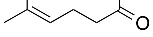
**10**
6-methyl-5-hepten-2-one
$C_8H_{14}O$ 126.10

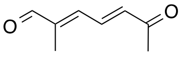
**11**
2-Methyl-6-oxo-2,4-heptadienal
$C_8H_{10}O_2$ 138.07

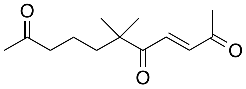
**12**
6,6-Dimethyl-3-undecene-2,5,10-trione
$C_{13}H_{20}O_3$ 224.14

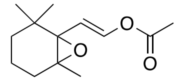
**13**
2-{2,2,6-Trimethyl-7-oxabicyclo[4.1.0]heptan-1-yl}ethenyl acetate
$C_{13}H_{20}O_3$ 224.14

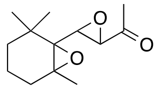
**14**
ß-Ionone-5,6-7,8-diepoxide
$C_{13}H_{20}O_3$ 224.14

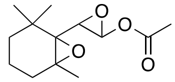
**15**
3-{2,2,6-Trimethyl-7-oxabicyclo[4.1.0]heptan-1-yl}oxiran-2-yl acetate
$C_{13}H_{20}O_4$ 240.14

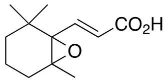
**16**
(2E)-3-{2,2,6-Trimethyl-7-oxabicyclo[4.1.0]heptan-1-yl}prop-2-enoic acid
$C_{12}H_{18}O_3$ 210.13

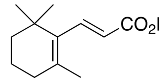
**17**
(2E)-3-(2,6,6-Trimethylcyclohex-1-en-1-yl)prop-2-enoic acid
$C_{12}H_{18}O_2$ 194.13

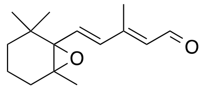
**18**
ß-Ionylidene acetaldehyde-5,6-epoxide
$C_{15}H_{22}O_2$ 234.16

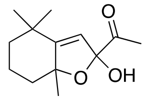
**19**
2-Hydroxy-4,4,7a-trimethyl-octahydro-1-benzofuran-2-yl acetate
$C_{13}H_{20}O_3$ 224.14

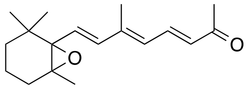
**20**
ß-Apo-13-carotenone-5,6-epoxide
$C_{18}H_{26}O_2$ 274.19

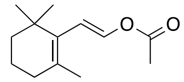
**21**
2-(2,6,6-Trimethylcyclohex-1-en-1-yl)ethenyl acetate
$C_{13}H_{20}O_2$ 208.15

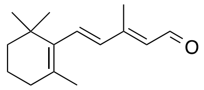
**22**
ß-Ionylidene acetaldehyde
$C_{15}H_{22}O$ 218.17

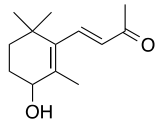
**23**
4-Hydroxy-ß-ionone
$C_{13}H_{20}O_2$ 208.15

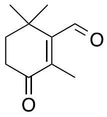
**24**
4-Oxo-ß-cyclocitral
$C_{10}H_{14}O_2$ 166.10

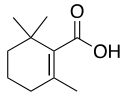
**25**
2,6,6-trimethyl-1-cyclohexene-1-carboxylic acid
$C_{10}H_{16}O_2$ 168.12

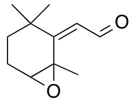
**26**
2-[(2Z)-1,3,3-Trimethyl-7-oxabicyclo[4.1.0]heptan-2-ylidene]acetaldehyde
$C_{11}H_{16}O_2$ 180.12

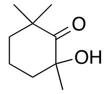
**27**
2-Hydroxy-2,6,6-trimethylcyclohexanone
$C_9H_{16}O_2$ 156.12

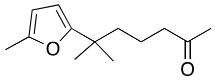
**28**
6-Methyl-6-(5-methylfuran-2-yl)heptan-2-one
$C_{13}H_{20}O_2$ 208.15

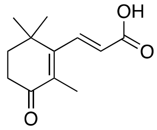
**29**
(2E)-3-(2,6,6-trimethyl-3-oxo-1-cyclohexen-1-yl)-2-propenoic acid
$C_{12}H_{16}O_3$ 208.11

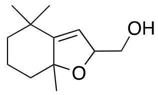
**30**
(4,4,7a-Trimethyl-2,4,5,6,7,7a-hexahydro-1-benzofuran-2-yl)methanol
$C_{12}H_{20}O_2$ 196.15

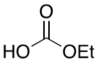
**31**
Ethyl bicarbonate
$C_3H_6O_3$ 90.03

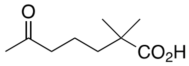
**32**
Geronic acid
$C_9H_{16}O_3$ 172.11
