## Supplemental File S2 for "Safety and uptake of fully oxidized β-carotene"

**Supplementary Material S2**. Average concentrations (µg/g or µg/mL) of polymer (Table 1) and apocarotenoids (Table 2) in tissues and fluids of mice dosed orally with a single dose of OxBC (300 mg/kg body weight) daily for 2 days and 5 days vs. control mice dosed with vehicle alone. The results were obtained from calibrated Multi-Reaction Monitoring (MRM) of data for the two transitions at 181→107 and 181→135 of DHA released from isolated OxBC polymer after NaOH treatment (Table 1) or present in OxBC apoC hexane extract (Table 2).

**Table 1**. OxBC polymer concentrations (µg/g or µg/mL) in tissues and body fluids.[1]

| | **Control** | | | | | **OxBC dose** | | | | |
|---|---|---|---|---|---|---|---|---|---|---|
| | **Day** | **Mouse** | **181→107** | **181→135** | **Average** | **Day** | **Mouse** | **181→107** | **181→135** | **Average** |
| **Kidney** | 2 | 1 | 0.96 | 1.3 | **1.1** | 2 | 7 | 2.2 | 2.8 | **2.5** |
| | 2 | 2 | 1.4 | 1.6 | **1.5** | 2 | 8 | 14 | 15 | **14** |
| | 5 | 4 | 1.8 | 2.2 | **2.0** | 5 | 10 | 4.0 | 5.0 | **4.5** |
| | 5 | 5 | 4.0 | 4.6 | **4.3** | 5 | 11 | 2.0 | 2.5 | **2.2** |
| | | | | *Average:* | ***2.2*** | | | | *Average:* | ***5.9*** |
| **Abdominal Muscle** | 2 | 1 | 3.1 | 3.3 | **3.2** | 2 | 7 | 2.2 | 2.8 | **2.5** |
| | 5 | 4 | 4.6 | 5.0 | **4.8** | 5 | 10 | 3.7 | 4.0 | **2.5** |
| | | | | *Average:* | ***4.0*** | | | | *Average:* | ***3.2*** |
| **Lung** | 2 | 3 | 13 | 20 | **16** | 2 | 8 | 19 | 22 | **21** |
| | 5 | 5 | 25 | 26 | **26** | 5 | 11 | 17 | 18 | **18** |
| | | | | *Average:* | ***21*** | | | | *Average:* | ***19*** |
| **Plasma** | 1 h | 5 | 0.20 | 0.24 | **0.22** | 1 h | 11 | 0.76 | 0.84 | **0.80** |
| | 1 h | 5 | 0.28 | 0.32 | **0.30** | 1 h | 11 | 1.4 | 1.5 | **1.42** |
| | 1 h | 6 | 0.55 | 0.76 | **0.65** | 1 h | 12 | 2.5 | 2.7 | **2.60** |
| | 2 | 2 | 0.09 | 0.05 | **0.07** | 2 | 8 | 0.90 | 0.95 | **0.93** |
| | 2 | 2 | 0.13 | 0.14 | **0.13** | 2 | 9 | 0.89 | 0.94 | **0.91** |
| | 2 | 3 | 0.17 | 0.29 | **0.23** | 5 | 11 | 0.76 | 0.77 | **0.76** |
| | 5 | 8 | 0.15 | 0.20 | **0.17** | 5 | 12 | 0.73 | 0.79 | **0.76** |
| | 5 | 5 | 0.22 | 0.25 | **0.23** | | | | *Average:* | ***1.2*** |
| | 5 | 10 | 0.27 | 0.43 | **0.35** | | | | | |
| | | | | *Average:* | ***0.26*** | | | | | |

| | | | | | | | | | | |
|---|---|---|---|---|---|---|---|---|---|---|
| **Stomach tissue** | 2 | 2 | 0.87 | 1.4 | **1.2** | 2 | 8 | 87 | 89 | **88** |
| | 2 | 3 | 22 | 23 | **23** | 2 | 7 | 265 | 267 | **266** |
| | 5 | 5 | 0.44 | 0.49 | **0.47** | 5 | 11 | 211 | 210 | **210** |
| | 5 | 4 | 12 | 11 | **11** | 5 | 10 | 844 | 826 | **835** |
| | | | | *Average:* | ***8.9*** | | | | *Average:* | ***350*** |
| **Feces** | 2 | 1 | 13.38 | 14.07 | **14** | 2 | 7 | 48.96 | 49.72 | **49.3** |
| | 2 | 3 | 7.94 | 8.17 | **8.1** | 2 | 9 | 123.28 | 125.68 | **124.5** |
| | 5 | 4 | 9.42 | 10.10 | **9.8** | 5 | 10 | 19.04 | 19.38 | **19.2** |
| | 5 | 6 | 11.94 | 14.39 | **13** | 5 | 12 | 48.49 | 52.93 | **50.7** |
| | | | | *Average:* | ***11*** | | | | *Average:* | ***61*** |
| **Intestinal wash** | 2 | 1 | 0.16 | 0.18 | **0.17** | 2 | 7 | 29.38 | 29.07 | **29** |
| | 5 | 4 | 0.23 | 0.24 | **0.24** | 5 | 10 | 3.31 | 3.31 | **3.3** |
| | | | | *Average:* | ***0.20*** | | | | *Average:* | ***16*** |
| **Urine** | 2 | 1 | 0.45 | 0.43 | **0.44** | 2 | 7 | 36.30 | 35.93 | **36** |
| | 2 | 3 | 0.98 | 0.72 | **0.85** | 2 | 9 | 61.56 | 62.46 | **62** |
| | 5 | 4 | 0.83 | 0.82 | **0.82** | 5 | 10 | 0.88 | 0.67 | **0.77** |
| | 5 | 6 | 1.35 | 1.60 | **1.48** | 5 | 12 | 29.25 | 30.30 | **30** |
| | | | | *Average:* | ***0.90*** | | | | *Average:* | ***32*** |
| **Cecum** | 2 | 1 | 6.96 | 8.73 | **7.8** | 2 | 7 | 30.36 | 39.49 | **35** |
| | 5 | 4 | 4.28 | 3.78 | **4.0** | 5 | 10 | 7.92 | 6.05 | **7.0** |
| | | | | *Average:* | ***5.9*** | | | | *Average:* | ***21*** |
| **Spleen** | 2 | 1 | 2.65 | 3.98 | **3.3** | 2 | 7 | 16.44 | 14.33 | **15** |
| | 5 | 4 | 3.92 | 4.84 | **4.4** | 5 | 10 | 12.70 | 12.01 | **12** |
| | | | | | ***3.8*** | | | | *Average:* | ***14*** |
| **Hamstring** | 2 | 1 | 2.5 | 3.2 | **2.8** | 2 | 7 | 2.0 | 1.9 | **2.0** |
| | 5 | 4 | 1.5 | 2.6 | **2.1** | 5 | 10 | 2.8 | 4.2 | **3.5** |
| | | | | *Average:* | ***2.5*** | | | | *Average:* | ***2.7*** |
| **Liver** | 2 | 3 | 113 | 113 | **113** | 2 | 9 | 81 | 80 | **80** |
| | 2 | 2 | 82 | 96 | **89** | 2 | 7 | 37 | 42 | **39** |

| | | | | | | | | | | |
|---|---|---|---|---|---|---|---|---|---|---|
| | 5 | 4 | 54 | 54 | **54** | 5 | 12 | 76 | 76 | **76** |
| | 5 | 4 | 30 | 33 | **31** | 5 | 10 | 86 | 87 | **86** |
| | 5 | 5 | 29 | 33 | **31** | | | | *Average:* | ***70.6*** |
| | | | | *Average:* | ***64*** | | | | | |
| **Small intestine tissue** | 2 | 1 | 6.2 | 7.1 | **6.6** | 2 | 7 | 4.1 | 4.0 | **4.1** |
| | 2 | 2 | 3.5 | 3.7 | **3.6** | 2 | 8 | 17 | 19 | **18** |
| | 5 | 4 | 1.1 | 1.6 | **1.4** | 2 | 9 | 24 | 26 | **25** |
| | 5 | 5 | 1.7 | 2.3 | **2.0** | 5 | 10 | 1.6 | 1.4 | **1.5** |
| | | | | *Average:* | ***3.4*** | 5 | 11 | 14 | 15 | **14** |
| | | | | | | 5 | 12 | 2.5 | 3.7 | **3.1** |
| | | | | | | | | | *Average:* | ***11.0*** |
| **Large intestine tissue** | 2 | 1 | 4.7 | 5.3 | **5.0** | 2 | 8 | 9.7 | 12 | **11** |
| | 2 | 2 | 4.1 | 6.0 | **5.0** | 2 | 9 | 6.1 | 6.6 | **6.4** |
| | 5 | 4 | 3.8 | 3.9 | **3.8** | 2 | 7 | 6.2 | 7.1 | **6.6** |
| | 5 | 5 | 1.6 | 2.6 | **2.1** | 5 | 11 | 3.9 | 4.2 | **4.1** |
| | | | | *Average:* | ***4.0*** | 5 | 12 | 3.9 | 4.2 | **4.0** |
| | | | | | | 5 | 10 | 3.5 | 3.7 | **3.6** |
| | | | | | | | | | *Average:* | ***5.9*** |

[1] Calibration was performed against OxBC polymer using DHA released by NaOH treatment

**Table 2**. Total apocarotenoid concentration (μg/g or μg/mL) of OxBC apoC in tissues and body fluids.[1]

| | Control | | | | | OxBC dose | | | | |
|---|---|---|---|---|---|---|---|---|---|---|
| | **Day** | **Mouse** | **181→107** | **181→135** | **Average** | **Day** | **Mouse** | **181→107** | **181→135** | **Average** |
| **Kidney** | 2 | 1 | 1.6 | 1.8 | **1.7** | 2 | 7 | 3.2 | 3.4 | **3.3** |
| | 2 | 2 | 1.9 | 2.0 | **2.0** | 2 | 8 | 3.6 | 3.9 | **3.8** |
| | 5 | 4 | 3.2 | 3.5 | **3.3** | 5 | 10 | - | - | - |
| | 5 | 5 | 1.5 | 1.5 | **1.5** | 5 | 11 | 2.4 | 2.5 | **2.5** |
| | | | | *Average:* | ***2.1*** | | | | *Average:* | ***3.2*** |
| **Abdominal muscle** | 2 | 1 | 2.6 | 2.5 | **2.6** | 2 | 7 | 3.0 | 3.0 | **3.0** |
| | 5 | 4 | 7.1 | 6.8 | **7.0** | 5 | 10 | 3.4 | 3.7 | **3.5** |
| | | | | *Average:* | ***4.8*** | | | | *Average:* | ***3.3*** |
| **Lung** | 2 | 3 | 10 | 10 | **10** | 2 | 8 | 21 | 22 | **21** |
| | 5 | 5 | 24 | 24 | **24** | 5 | 11 | 6 | 4 | **5** |
| | | | | *Average:* | ***17*** | | | | *Average:* | ***13*** |
| **Plasma** | 1 h | 5 | 0.004 | 0.004 | **0.004** | 1 h | 11 | 0.002 | 0.002 | **0.002** |
| | 1 h | 6 | 0.109 | 0.112 | **0.11** | 1 h | 12 | 0.28 | 0.28 | **0.28** |
| | 2 | 2 | 0.001 | 0.001 | **0.001** | 2 | 8 | 0.11 | 0.08 | **0.10** |
| | 2 | 3 | 0.073 | 0.064 | **0.069** | 2 | 9 | 0.12 | 0.14 | **0.13** |
| | 5 | 8 | 0.001 | 0.001 | **0.001** | 5 | 12 | 0.09 | 0.08 | **0.08** |
| | 5 | 5 | 0.024 | 0.036 | **0.030** | | | | *Average:* | ***0.12*** |
| | 5 | 10 | 0.048 | 0.055 | **0.051** | | | | | |
| | | | | *Average:* | ***0.038*** | | | | | |
| **Stomach** | 2 | 2 | 1.3 | 1.3 | **1.3** | 2 | 8 | 38 | 38 | **38** |
| | 2 | 3 | 2.1 | 2.1 | **2.1** | 2 | 7 | 37 | 37 | **37** |
| | 5 | 5 | 1.0 | 1.0 | **1.0** | 5 | 11 | 80 | 80 | **80** |
| | 5 | 4 | 2.6 | 2.6 | **3** | 5 | 10 | 138 | 138 | **138** |

| | | | | | | | | | | |
|---|---|---|---|---|---|---|---|---|---|---|
| | | | | *Average:* | ***1.7*** | | | | *Average:* | ***73*** |
| **Feces** | 2 | 1 | 0.50 | 0.54 | **0.52** | 2 | 7 | 1.6 | 2.4 | **2.0** |
| | 2 | 3 | 0.38 | 0.33 | **0.36** | 2 | 9 | 2.4 | 2.1 | **2.2** |
| | 5 | 4 | 0.50 | 0.50 | **0.50** | 5 | 10 | 1.1 | 1.2 | **1.2** |
| | 5 | 6 | 0.52 | 0.61 | **0.57** | 5 | 12 | 2.1 | 2.3 | **2.2** |
| | | | | *Average:* | ***0.49*** | | | | *Average:* | ***1.9*** |
| **Intestinal wash** | 2 | 1 | 0.009 | 0.015 | **0.012** | 2 | 7 | 0.31 | 0.36 | **0.34** |
| | 5 | 4 | 0.010 | 0.014 | **0.012** | 5 | 10 | 0.11 | 0.10 | **0.10** |
| | | | | *Average:* | ***0.012*** | | | | *Average:* | ***0.22*** |
| **Urine** | 2 | 1 | 0.12 | 0.10 | **0.11** | 2 | 7 | 0.76 | 0.94 | **0.85** |
| | 2 | 3 | 0.25 | 0.36 | **0.31** | 2 | 9 | 0.61 | 0.64 | **0.62** |
| | 5 | 4 | 0.52 | 0.78 | **0.65** | 5 | 10 | 0.45 | 1.21 | **0.83** |
| | 5 | 6 | 0.15 | 0.19 | **0.17** | 5 | 12 | 0.21 | 0.25 | **0.23** |
| | | | | *Average:* | ***0.31*** | | | | *Average:* | ***0.63*** |
| **Cecum** | 2 | 1 | 0.44 | 0.49 | **0.46** | 2 | 7 | 0.72 | 0.84 | **0.78** |
| | 5 | 4 | 0.37 | 0.42 | **0.40** | 5 | 10 | 1.3 | 1.5 | **1.4** |
| | | | | *Average:* | ***0.43*** | | | | *Average:* | ***1.1*** |
| **Spleen** | 2 | 1 | 0.10 | 0.14 | **0.12** | 2 | 7 | 0.10 | 0.14 | **0.12** |
| | 5 | 4 | 0.11 | 0.13 | **0.12** | 5 | 10 | 0.32 | | **0.32** |
| | | | | *Average:* | ***0.12*** | | | | *Average:* | ***0.22*** |
| **Hamstring** | 2 | 1 | 23 | 23 | **23** | 2 | 7 | 16 | 16 | **16** |
| | 5 | 4 | 11 | 11 | **11** | 5 | 10 | 15 | 15 | **15** |
| | | | | *Average:* | ***17*** | | | | *Average:* | ***16*** |
| **Liver** | 2 | 3 | 3.6 | 3.6 | **3.6** | 2 | 9 | 2.6 | 2.5 | **2.6** |
| | 2 | 2 | 3.6 | 3.6 | **3.6** | 2 | 7 | 3.5 | 3.5 | **3.5** |
| | 5 | 4 | 2.5 | 2.5 | **2.5** | 5 | 12 | 3.5 | 3.5 | **3.5** |
| | 5 | 4 | 2.6 | 2.5 | **2.6** | 5 | 10 | 4.8 | 4.8 | **4.8** |
| | 5 | 5 | 2.5 | 2.5 | **2.5** | | | | *Average:* | ***3.6*** |

| | | | | | | | | | | |
|---|---|---|---|---|---|---|---|---|---|---|
| | | | | *Average:* | ***3.0*** | | | | | |
| **Small intestine tissue** | 2 | 1 | 0.29 | 0.30 | **0.29** | 2 | 8 | 0.72 | 0.67 | **0.69** |
| | 2 | 2 | 0.17 | 0.16 | **0.17** | 2 | 9 | 1.0 | 1.0 | **1.0** |
| | 5 | 4 | 0.15 | 0.15 | **0.15** | 5 | 11 | 1.2 | 1.3 | **1.2** |
| | 5 | 5 | 0.24 | 0.24 | **0.24** | 5 | 12 | 0.22 | 0.23 | **0.22** |
| | | | | *Average:* | ***0.21*** | | | | *Average:* | ***0.79*** |
| **Large intestine tissue** | 2 | 1 | 0.81 | 0.05 | **0.43** | 2 | 8 | 0.46 | 0.54 | **0.50** |
| | 2 | 2 | 0.12 | 0.14 | **0.13** | 2 | 9 | trace | trace | **trace** |
| | 5 | 4 | trace | trace | **trace** | 5 | 11 | 0.18 | 0.19 | **0.18** |
| | 5 | 5 | trace | trace | **trace** | 5 | 12 | 0.22 | 0.24 | **0.23** |
| | | | | *Average:* | ***0.28*** | | | | *Average:* | ***0.30*** |

[1] Calibration was performed against OxBC using DHA present in hexane extract of OxBC
